# Unconscious manipulation of conceptual representations with decoded neurofeedback impacts search behaviour

**DOI:** 10.1101/2023.07.04.547632

**Authors:** Pedro Margolles, Patxi Elosegi, Ning Mei, David Soto

**Author notes:** Correspondence should be addressed to, Basque Center on Cognition, Brain and Language, Paseo Mikeletegi 69, 2nd Floor 20009 San Sebastian.

## Abstract

The necessity of conscious awareness in human learning has been a long-standing topic in psychology and neuroscience. Previous research on non-conscious associative learning is limited by the low signal-to-noise ratio of the subliminal stimulus, and the evidence remains controversial, including failures to replicate. Using functional MRI decoded neurofeedback (fMRI-DecNef) we guided participants from both sexes to generate neural patterns akin to those observed when visually perceiving real-world entities (e.g., dogs). Importantly, participants remained unaware of the actual content represented by these patterns. We utilized an associative DecNef approach to imbue perceptual meaning (e.g., dogs) into Japanese hiragana characters that held no inherent meaning for our participants, bypassing a conscious link between the characters and the dogs concept. Despite their lack of awareness regarding the neurofeedback objective, participants successfully learned to activate the target perceptual representations in the bilateral fusiform. The behavioural significance of our training was evaluated in a visual search task. DecNef and control participants searched for dogs or scissors targets that were pre-cued by the hiragana used during DecNef training or by a control hiragana. The DecNef hiragana did not prime search for its associated target but, strikingly, participants were impaired at searching for the targeted perceptual category. Hence, conscious awareness may function to support higher-order associative learning. Meanwhile, lower-level forms of re-learning, modification, or plasticity in existing neural representations can occur unconsciously, with behavioural consequences outside the original training context. The work also provides an account of DecNef effects in terms of neural representational drift.

**Significance Statement:** This study examined the role of conscious awareness in human learning by using fMRI-DecNef. These techniques enabled participants to self-regulate their brain activity to align with the perceptual representations generated by a real-world entity (i.e., dogs), without awareness of the content they represented. We demonstrated that established brain conceptual representations can be unconsciously modified, influencing visual search behaviour for the targeted perceptual content through the neural representational drift mechanism. Nonetheless, our research suggests that conscious awareness plays a role in more advanced forms of associative learning. Further, this study offers methodological insights for improving DecNef protocols and suggests potential for personalized interventions, including guidance to correct maladaptive conceptual representations.

## Introduction

The acquisition of word meanings depends on associative learning of symbols and episodic referents (Blewitt, 1982; M. D. Smith, 1978). A longstanding question in psychology and neuroscience concerns whether conscious awareness is necessary for higher-level associative learning.

Prior research provided evidence of unconscious associative learning between subliminal visual items and conscious rewards (Pessiglione et al., 2008; Seitz, Kim, & Watanabe, 2009; Skora, Yeomans, Crombag, & Scott, 2021), as well as fear conditioning responses to stimuli not consciously perceived (Raio, Carmel, Carrasco, & Phelps, 2012). Studies using evaluative learning paradigms have suggested that emotional value can be conditioned into neutral visuals, like nonsense syllables, and that these associations can be made implicitly, even without participants awareness (Hofmann, De Houwer, Perugini, Baeyens, & Crombez, 2010; Staats & Staats, 1957; Tryon & Cicero, 1989). However, the aforementioned evidence is controversial and has faced substantial criticism (Field, 2000; Mertens & Engelhard, 2020; Shanks & Dickinson, 1990; Shanks & John, 1994), raising the possibility that the results were due to experimental demand characteristics and participants’ awareness of the experimental goals, not from non-conscious knowledge (Field, 2000; Page, 1974; Stahl, Haaf, & Corneille, 2016).

Research on unconscious associative learning is limited due to the low signal-to-noise ratio of the brief and masked unconscious stimulus. Learning may falter in these conditions. Addressing this challenge requires an increase in signal strength in the unconscious condition (Lau, 2022). Here, we used fMRI-DecNef to enable participants generate brain patterns matching a target state without awareness of the content represented by those brain patterns (Watanabe, Sasaki, Shibata, & Kawato, 2017). Previous DecNef studies mainly centered on modulating affective responses to faces (Shibata, Watanabe, Kawato, & Sasaki, 2016), fear (Koizumi et al., 2016; Taschereau-Dumouchel et al., 2018) or confidence (Cortese, Amano, Koizumi, Kawato, & Lau, 2016). Here we used DecNef training to shape brain representations and form novel conceptual knowledge. Specifically, we aimed to incept the perceptual meaning of a real-world entity into a Japanese hiragana that had no inherent meaning for our participants, by reinforcing related neural patterns. To achieve this, we developed an open framework for DecNef that integrates several enhancements to the standard pipeline, namely, a customised decoder that provides more fine-grained and less binomial/overconfident predictions during DecNef training (Margolles, Mei, Elosegi, & Soto, 2023).

First, we trained a classifier to differentiate between brain patterns associated with perceiving dogs or scissors images (Figure 1A). During several DecNef training sessions, participants self-regulated their brain activity while viewing a hiragana to maximize the size of a feedback disk and monetary reward (Figure 1B). Participants were unaware that the feedback depended on how closely their brain activity patterns matched the classifier’s representation of dogs in the bilateral fusiform. We chose the fusiform due to its role in object recognition and imagery (Dijkstra, van Gaal, Geerligs, Bosch, & van Gerven, 2021; Lee, Kravitz, & Baker, 2012; Reddy, Tsuchiya, & Serre, 2010; Stokes, Thompson, Cusack, & Duncan, 2009). We anticipated that this approach would create associative knowledge, imbuing the hiragana with the meaning of dogs. The behavioural significance of the DecNef was evaluated using a visual search task involving dogs and scissors, with the search target being pre-cued by the DecNef hiragana or by a control hiragana (Figure 1C). We hypothesized that associating the hiragana to the dogs concept would lead to automatic attention shifts towards the matching search item, resulting in faster performance and distinct neural responses compared to the control hiragana (Soto, Hodsoll, Rotshtein, & Humphreys, 2008; Soto, Humphreys, & Rotshtein, 2007). Additionally, we compared the search behavior of the DecNef group to a control group to understand better how DecNef training influenced the target category’s representation itself. This contrast explores the potential of DecNef training to unconsciously modify the neural representation of the target concept, resulting in distinct search performance for dogs targets compared to scissors.

**Figure 1:**
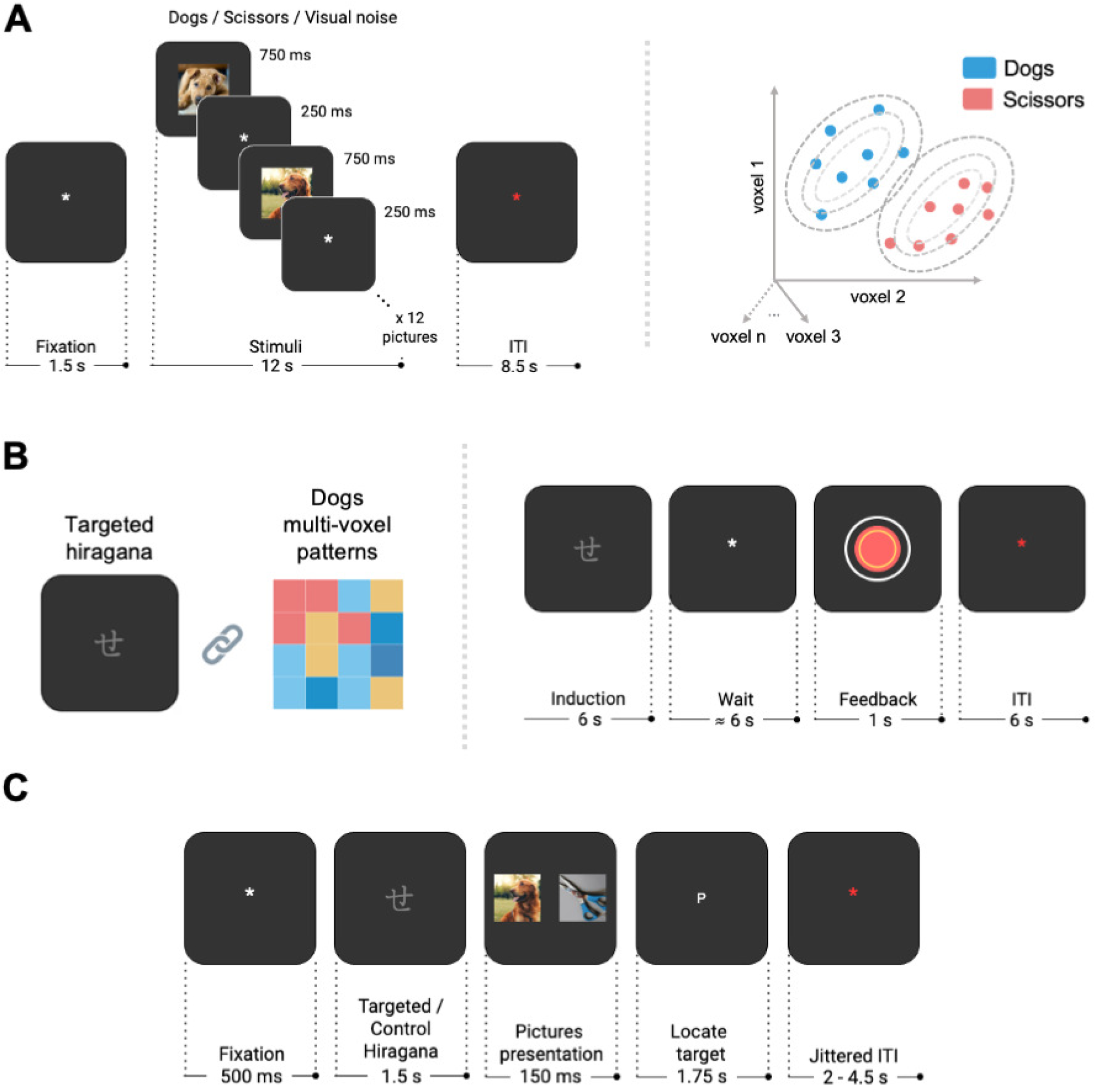
Overview of the experimental design. (A) Participants viewed pictures of dogs and scissors. We trained a custom logistic regression decoder using perceptual data from the bilateral fusiform to distinguish between the multivoxel patterns associated with each class. (B) Over four DecNef sessions, our goal was to covertly associate a neutral hiragana with the perceptual meaning of dogs using reinforcement learning. Participants self-regulated their brain activity in the bilateral fusiform while looking at an hiragana. Monetary rewards were given based on the alignment of their brain activity patterns with the target class (i.e., dogs) as determined by the classifier. The decoded volumes matched the three fMRI volumes acquired between 5 and 11 seconds after hiragana onset, aligning with the peak of the Hemodynamic Response Function during self-regulation. Importantly, participants remained unaware of the experiment’s true objective and the existence of a target category. (C) Upon completion of the four DecNef sessions, a visual search task was administered to evaluate the behavioural and neural impact of the DecNef training. Participants looked for dogs or scissors targets, which were either pre-signaled by the hiragana used during DecNef training or by a new hiragana. We anticipated changes in search times due to the unconscious priming of dogs’ perceptual representations driven by the targeted hiragana.

## Materials and Methods

### Participants

The DecNef training involved a group of 18 young adults (23-36 years old, mean*±*SD age: 28.19*±*3.73 years, 6 males) with normal or corrected-to-normal vision, who were right-handed and unfamiliar with the Japanese language. A sample size of 18 is well above average in DecNef studies (Cortese et al., 2016; Koizumi et al., 2016; Taschereau-Dumouchel et al., 2018). The experimental procedures were conducted in accordance with the Declaration of Helsinki and approved by the BCBL Research Ethics Board. All participants provided written informed consent and at the end of the experiment, received a fixed monetary compensation of 105€ for their participation in the study. Additionally, we included a control group of 36 young participants to double the DecNef group sample size and improve the statistical power of our analysis for detecting any differences between groups (20-29 years old, mean*±*SD age: 23.16*±*2.62 years, 11 males). One of the participants had to be replaced due to a high number of errors (i.e., > 40%).

We also note here that the decoded neurofeedback paradigm is a special case when it comes to the selection of a control group. Standard fMRI neurofeedback studies are based on explicit approaches in which participants are aware of the strategy that should be used to modulate brain activity in the targeted region, and hence standard fMRI neurofeedback approaches use control groups in which participants receive feedback of a non-target brain region, sham feedback, or perform mental strategies without neurofeedback signaling (Sorger, Scharnowski, Linden, Hampson, & Young, 2019). In the case of DecNef, and specifically in our associative DecNef study, the above is not applicable (Sorger et al., 2019) because the neurofeedback training occurs at an implicit level, namely, participants are unaware of (i) the target brain region (ii) what the neurofeedback signal represents (iii) the strategy that may be used to regulate activity in the target region. Because of this, we elected to include a control group of participants that did not undergo DecNef in order to test that the behavioral pattern of search performance in the DecNef group was indeed specific to DecNef training. Note that a similar approach was used in prior associative DecNef studies (Amano, Shibata, Kawato, Sasaki, & Watanabe, 2016; Knotts, Cortese, Tascherau-Dumouchel, Kawato, & Lau, 2019).

### Apparatus

Our DecNef pipeline required three high-performance computers: the computer controlling the MRI scanner; a server computer, which analyzed the data in real time; and a client computer for feedback and stimuli presentation during MRI scanning. The three computers were connected through an ethernet local area network.

### MRI settings

Whole-brain data was acquired on a 3T Siemens Magnetom Prisma-fit whole-body MRI scanner with a 64-channel head-coil at the BCBL facilities. A gradient-echo echo-planar imaging sequence was used with the following parameters: time-to-repetition (TR) = 2000 ms, time-to-echo (TE) = 28 ms, flip angle (FA) = 74°, field of view (FoV) = 192 mm, 34 axial slices with no inter-slice gap and a 3*3*3.5 mm^3^ voxel resolution. Five volumes were acquired as MRI heat-up at the beginning of each fMRI run to ensure steady-state magnetization. These volumes were then discarded from analyses. Additionally, one high-resolution T1-weighted structural MPRAGE scan was acquired: TR = 2530 ms, TE = 2.36 ms, FA = 7°, FoV = 256 mm, 1 mm isotropic voxel resolution, 176 slices. Visual stimuli were presented on an out-of-bore screen, which participants viewed through a mirror mounted on the head coil. To minimize head movements, participants were instructed to keep their head as still as possible, and the head-coil of the MRI scanner was cushioned with foam. Participants used padded headphones to minimize background scanner noise, to facilitate the presentation of auditory stimuli, and to enable communication with experimenters via a compatible microphone.

### fMRI scans for decoder construction

The decoder construction session involved four perceptual runs and two mental imagery runs, each lasting approximately 8 minutes.

First, participants completed four perceptual runs in which they were presented with pictures of dogs and scissors from the THINGS database (Hebart et al., 2019). Each perceptual run was composed of seven dogs and seven scissors trials, and every trial had a duration of 22 seconds. A trial started with a white fixation point lasting 1500 ms (.37 *^◦^* of visual angle), positioned at the center of the screen against a black background. This was followed by the sequential presentation of 12 example pictures (8.06*^◦^*) from a specific visual category (dogs or scissors), with each image lasting for 750 ms each. Picture presentation were interleaved with 250 ms visual fixation periods. The same images were used for each category across trials, but were presented in random order. Finally, to maximize the separation between the brain activity patterns across trials, we included a 8500 ms inter-trial interval (ITI) with a red fixation (.37 *^◦^*) at the center of the screen (Figure 1A). During the perceptual runs, participants were instructed to focus their attention on the presented pictures and actively engage in processing their meaning. Additionally, each perceptual run featured four trials where visual-noise patterns were displayed. These gaussian noise patterns were crafted by randomly selecting colored pixels with varying intensities from the RGB (Red, Green, Blue) spectrum. In each trial, 12 of these patterns were shown (8.06 *^◦^*), chosen at random from a pool of 60 samples. The data gathered from the visual noise trials were also used to train the decoder, allowing us to tackle representational noise when distinguishing between dogs and scissors perceptual representations during DecNef training (see the ‘Algorithm for decoding’ section for more details).

Furthermore, we sought to consider that any representation induced during DecNef, resembling the target representations, would be internally generated. Consequently, following the perceptual runs, participants underwent two mental imagery runs. During these runs, they were instructed to vividly imagine a picture of their choice from each of the visual categories. This image was chosen at the start of the perceptual runs based on its ease of imagination for the participant. Each mental imagery run comprised nine imagery trials for every category, with each trial lasting 22 seconds. These trials began with a 1500 ms white fixation (.37*^◦^*) against a black background, succeeded by a 12-second period where participants were guided to imagine the chosen category image, all while holding their fixation. To indicate which category they should imagine, auditory prompts in Spanish, either ‘Perro’ (meaning dog; 375 ms) or ‘Tijeras’ (meaning scissors; 676 ms), were played three times during the 12-second span. Auditory cues were used to avoid any visual stimulation that could interfere with visual imagery. The volume of the auditory cues was adjusted individually for each participant. After the onset of the auditory prompt, participants had to visualize for approximately 4.5 seconds until the next auditory cue began. They were instructed to mentally project a vivid and static image at the center of the screen. Upon hearing the second and third auditory cues, participants were directed to re-activate the target image for that trial. Between trials, an ITI with a red fixation point (.37 *^◦^*) was displayed at the screen’s center for 8500 ms. Following both the perceptual and mental imagery runs, participants were invited to take a short break of 1-2 minutes. We used OpenSesame software (Mathôt & March, 2022; Mathôt, Schreij, & Theeuwes, 2012) to control stimulus presentation.

### fMRI training data preprocessing

The fMRI volumes used for training the decoder were preprocessed using a customized pipeline in AFNI v21.0.05 (Cox, 1996), which was created through the Nipype v.1.4.1 (Gorgolewski et al., 2011) interface module for Python 3.6.

The first experimental volume from the decoder construction session, obtained after the heat-up volumes, served as a functional reference for the realignment of volumes for each participant. That reference DICOM file was converted to NifTi format using Dcm2Niix (Li, Morgan, Ashburner, Smith, & Rorden, 2016) and de-obliqued to match cardinal coordinate orientation with AFNI 3dWarp command.

The subsequent perceptual and mental imagery volumes were co-registered to the reference volume using AFNI 3dvolreg command using heptic (7th order) Lagrange polynomial interpolation (Oakes et al., 2005). The OpenSesame task logs were used to label all volumes, indicating their corresponding stimulus category, trials onset times, and fMRI run. For each run, we linearly detrended the fMRI time series to remove drift before training the decoder. To match the characteristics of the Hemodynamic Response Function (HRF) and capture the peak neural activity associated with the presented stimuli, we selected these fMRI volumes occurring between 5 and 18 seconds following the initial picture/auditory prompt presentation. This spanned 7 volumes per trial in both the perception and mental imagery runs. The selected perceptual and mental imagery samples were stacked independently for ROI delineation and decoder training. Next, the stacked perceptual volumes were z-scored by subtracting the samples’ mean value at each voxel and dividing by the standard deviation. The standardization of the scans in the imagery runs was performed by using the voxels mean and standard deviation from the perceptual samples. This approach was utilized to establish a uniform and comparable scale for the data from both conditions, and investigate the extent to which information could generalize from perception to mental imagery.

### Algorithm for decoding

To establish the ROI for DecNef training, our initial step was to outline the regions where perceptual representations, triggered by the pictures, converged with internally generated representations during mental imagery. Previous research had identified these overlapping regions in the fusiform gyrus (Dijkstra et al., 2021; Lee et al., 2012; Reddy et al., 2010; Stokes et al., 2009). The rationale behind this approach is that the brain activity patterns used for training the classifier are mainly externally triggered through the presentation of the images, while the relevant brain activity patterns during DecNef training may stem from internal mental simulation. Therefore, our goal was to enhance the cross-domain generalization of our machine learning classifier and facilitate DecNef training by focusing on these voxels located at the overlap between perception and imagery in the fusiform region.

Accordingly, a searchlight decoding analysis was conducted for each participant. The initial step involved creating a binarized functional brain mask from the skull-stripped reference brain volume. Next, z-scored volumes from the perceptual and mental imagery runs were masked to exclude non-brain voxels with zero values. Then, we performed a cross-domain searchlight analysis in the subject’s native space using Brain Imaging Analysis Kit (BrainIAK v. 0.9.1; http://brainiak.org; (Kumar et al., 2021). A searchlight of 9 * 9 * 9 voxels was defined by a cube cluster of 4-voxel (3 mm * 4 voxels = 12 mm) radius. We employed the scans from the perceptual runs, excluding data from visual noise samples, to train a standard logistic regression classifier with L2 regularization within each searchlight. This classifier was then applied to the imagery data in order to decode the content of mental imagery volumes as either dogs or scissors categories. We evaluated the decoder’s performance by computing the area under the receiver operating characteristic curve (AUC) and subsequently assigned this performance score to the center voxel of each searchlight. This resulted in a comprehensive whole-brain map of cross-domain decoding results.

Following this, we manually generated a spherical mask in each participant’s original functional space. The center of this mask was positioned around the voxels demonstrating the highest level of generalization from perception to mental imagery. In doing so, we analyzed the resulting searchlight maps across various mental imagery maps to draw our fusiform ROIs. The goal was to guarantee that the ultimate ROI for DecNef encompassed areas where the overlap between perception and imagery remained consistent over time. Visualizing the searchlight maps for each run allowed us to achieve a balance between selecting an overly narrow ROI and an overly broad ROI. Accordingly, this approach serves to address potential issues, such as neglecting spatial variability during DecNef training due to a narrow ROI or the risk of overfitting our decoder because of an excessively broad ROI. Further, the fusiform gyrus encompasses various regions with distinct functional and representational roles (see Pitcher, Charles, Devlin, Walsh, and Duchaine, 2009; Schwarzlose, Baker, and Kanwisher, 2005 in relation to faces, bodies, and objects). Opting for selecting an overly broad ROI and overly distributed voxels across the fusiform would pose a challenge to interpreting the outcomes of DecNef training. Therefore, we chose to utilize searchlight maps as a method to pinpoint fusiform areas characterized by more consistent functional, spatial, and temporal patterns.

The brain areas showing better classification performance varied to some extent across participants. Although our a priori target was on the fusiform region, the first participant exhibited noteworthy crossdomain decoding, particularly in the parietal cortex. Consequently, we incorporated this region alongside the bilateral fusiform into the DecNef training. The second and third participants exhibited greater decoding performance in the left fusiform gyrus compared to the right, leading us to choose the left fusiform as the target for DecNef. However, for subsequent participants, we opted to utilize a uniform region for DecNef training, centered in the bilateral fusiform gyrus. Across these three participants, this region consistently demonstrated a high degree of decoding accuracy. We aimed to ensure uniform ROI dimensions for both the left and right fusiform areas across all participants. The masks, on average, covered 561.35 voxels per hemisphere, with a standard deviation of *±*150.29 voxels. The spheres encompassing the left and right fusiform regions were combined to form a single bilateral fusiform mask. The stacked perceptual and mental simulation data was ultimately masked to preserve only the voxels located within the targeted bilateral fusiform ROI.

Subsequently, for each participant, we trained a custom decoder on the ROI mask obtained through the aforementioned searchlight analysis. The perceptual data served as the training set, and the classifier hyperparameters were optimized using the mental imagery data as the validation dataset. This method enhanced the decoder’s ability to generalize to the DecNef training sessions. During the development of our DecNef pipeline, we observed that sparse logistic regression classifiers (Yamashita, Sato, Yoshioka, Tong, & Kamitani, 2008), as commonly employed in previous DecNef training studies (Cortese et al., 2016; Cortese, Amano, Koizumi, Lau, & Kawato, 2017; Koizumi et al., 2016; Taschereau-Dumouchel et al., 2018), tended to produce highly bimodal/overconfident probability distributions for out-of-sample distribution predictions in the imagery or DecNef runs (Bai, Mei, Wang, & Xiong, 2021). Moreover, prior DecNef research did not explicitly address representational noise during DecNef training (Watanabe et al., 2017), which encompasses neural patterns that do not directly correspond to the specific trained representations or categories. This factor may reduce the capacity of the decoder to discriminate target from non-target representations, constraining the feedback signal and thereby hinder participants’ learning. These issues were addressed by developing a custom logistic regression classifier using Keras (Chollet, 2015) (v2.7) and Tensorflow (Abadi et al., 2016) (v2.7). The classifier consisted of a fully connected input and a softmax output layer, specifically designed for the binary classification of samples into dogs or scissors categories. To prevent overfitting, we applied regularization to the kernel weights and layer outputs using an L2 kernel regularizer of .1 and an L1 activity regularizer of .001, respectively. We used a default Stochastic Gradient Descent optimizer with a learning rate of .0001. Specifically, we employed Binary Cross-Entropy loss as the loss function with scissors labeled as 0, dogs as 1, and visual noise images as 0.5. By adopting this strategy, we achieved a more continuous representational gradient between categories and decreased the network’s susceptibility to the representational noise during testing. As a result, the network exhibited improved discrimination between the relevant classes by generating more moderated predicted probabilities. To evaluate the performance of our decoder, we utilized the AUC. We set a tolerance of .0001, which represents the minimum change in the validation loss required to qualify as an improvement. We also established a patience of 50 epochs; if there was no improvement in the validation loss after these 50 epochs, the training would be halted. Finally, we utilized a batch size of 64 to compute the gradient during each iteration. A comparison among the standard and sparse regression decoders, as well as our custom classifier using BCE Loss can be found in Margolles et al. (2023).

### DecNef training procedure

The DecNef training consisted of four fMRI sessions, each lasting approximately 70 minutes, performed on consecutive days at a rate of one session per day in the early morning (8am). In the first three sessions, there were a total of eight fMRI runs. These runs had a duration of approximately 6.5 minutes each and consisted of 18 trials per run, with each trial lasting approximately 19 seconds. In the final session, only two fMRI runs were carried out, both of the same duration as those in the earlier sessions. Subsequently, a visual search task was performed within the MRI scanner to serve as a behavioral assessment for DecNef training. A 2-minute rest period followed the completion of each run.

During the DecNef training sessions, participants encountered one of two gray Japanese hiragana characters (の or せ), with the presentation of each character being counter-balanced across participants to ensure equal distribution. Since the participants were not familiar with the Japanese language, these characters had no inherent meaning for them. Our goal was to establish a novel unconscious association between the perceptual representation of dogs and these neutral hiragana characters through reinforcement and associative learning.

Considering that distinct stimulus categories, such as living and non-living entities, are associated to distinct multivoxel patterns and neural substrates (Pitcher et al., 2009), attempting to counterbalance the target classes (i.e., dogs and scissors) would have led to the formation of two smaller groups, each comprising only nine subjects. In such circumstances, it becomes notably more difficult to derive meaningful conclusions or discern significant effects, largely due to the limited statistical power associated with smaller sample sizes. Opting to concentrate on a single category across all participants, would ensure a more robust evaluation of DecNef’s neural effects. Additionally, this approach would contribute to reducing potential sources of variability in participants’ behavioral responses. This approach has been utilized in previous associative DecNef studies (Amano et al., 2016; Knotts et al., 2019). Due to these considerations and the known responsiveness of the bilateral fusiform gyrus ROI to living entities (Connolly et al., 2012; Downing, Chan, Peelen, Dodds, & Kanwisher, 2006; Sha et al., 2015), indicating that targeting dogs representations might be more effective in inducing neural changes, we chose to assign dogs as the target category for all participants.

The entire experimental procedures were carried out implicitly. Participants were aware that they were participating in a neurofeedback study aimed at self-regulation of brain activity, but they were kept entirely unaware of the true purpose of the intervention, which included the existence of a target category. To prevent participants from associating the prior fMRI session used for the decoder construction with the subsequent DecNef training sessions, a new experimenter was assigned, and they were explicitly informed that they were taking part in a different experiment. Additionally, participants were asked to complete different consent forms and were given separate information sheets.

We instructed participants to engage in self-regulation of their brain activity with the goal of maximizing their monetary rewards by enlarging the size of a red circle displayed as feedback. At the beggining of the DecNef training, participants were told that they would receive a fixed compensation of 85€ for their participation in the experiment. Additionally, they had a chance to receive a performance-based bonus of €25, with the potential to earn up to €0.10 for each trial. Participants were explicitly encouraged to freely explore various mental strategies and choose those that proved most effective in achieving their goal. Nonetheless, we emphasized the importance of avoiding strategies that could introduce potential artifacts to the fMRI analysis, such as changes in breathing or body movements. After each session, participants were given an estimated report of their earnings designed to maintain their motivation. During the final DecNef session, participants engaged in only two fMRI runs and were encouraged to employ their most effective mental strategies from previous sessions to maximize the size of the red circle. However, in order to fulfill potential ethical considerations, all participants were given a fixed amount of 110€ at the end of the study, which included the 85€ compensation for their participation and the 25€ bonus reward.

At the outset of each trial (Figure 1B), the hiragana character was displayed on a black background for 6 seconds, subtending a visual angle of 1.29 *^◦^*. During its presentation, participants were directed to modulate their brain activity to enlarge a feedback disk that was to be presented subsequently. No further specific guidance or instructions were provided. Next, participants were given a 6-second relaxation period during which they were presented with a white fixation mark (.37 *^◦^*) while awaiting feedback. The feedback was then displayed as a red circle for one second. The circle’s size reflected the decoder’s probability of the target class, ranging from 0 to 1. This probability represented the similarity between the brain representations evoked during hiragana presentation and the brain activity patterns associated with dogs according to the previously trained decoder. We informed participants that their monetary reward would increase proportionally to the size of the red circle displayed as feedback. The maximum reward per trial (i.e., .1€) was indicated by a white border (3.67 *◦* of visual angle) enclosing the red circle, signifying 100% similarity between the participant’s brain activity and the decoder’s perceptual representations of dogs. Additionally, a yellow circumference was used to set a threshold below which no reward would be given for that trial. This threshold was set at 50% similarity. If participants reached a point where they were no longer making progress or their performance had plateaued, we encouraged them to assess their current mental strategies and experiment with new ones to achieve better outcomes. Each trial concluded with a fixed inter-trial interval of 6 seconds, during which a red fixation mark was displayed (.37 *◦*).

The fMRI scans during DecNef training were co-registered with the functional reference volume acquired during the decoder construction session, using the same preprocessing pipeline utilized for training the decoder. To ensure timely processing in real-time and prevent delays in analyzing incoming volumes, it was crucial that the co-registration times were consistently shorter than the fMRI TR. Here, co-registration of each volume took approximately 0.7 seconds (vs. TR = 2 seconds). Then, we applied the functional mask of the bilateral fusiform ROI extracted for each participant. Spatial smoothing was not performed. Linear detrending was applied to the ROI data, encompassing all scans available from a particular run up to the current volume, and including an extra set of 20 baseline scans collected at the onset of each run. Previous neurofeedback studies used z-scoring procedures based on run/session baselines or prior data from the current run (Oblak, Sulzer, & Lewis-Peacock, 2019). In contrast, we utilized the mean and standard deviation parameters for each voxel obtained from the scans used in decoder construction, specifically the perceptual data, for standardizing the volume data. This method does not impose any data leakage on the information that is used for decoding, because this procedure does not use any information regarding the classes. Moreover, the classification analysis is performed on the multivariate pattern across voxels and the voxelwise standardization procedure is blind to these patterns. Using the mean and standard deviation from the decoder construction data for voxel-wise standardization in the DecNef scans merely aligns the distributions of the train and test sets at the single voxel level, providing several benefits. For example, this approach enhances result interpretability by maintaining consistency across all runs and sessions with a common reference data distribution. Additionally, it ensures that the test data scales to match the distribution of the training data, potentially enhancing the decoder’s performance during DecNef training (see Margolles et al. 2023 for a comprehensive comparison of how different DecNef standardization techniques affect decoding performance). Real-time decoding focused on volumes captured between 5 and 11 seconds after the hiragana presentation, resulting in the decoding of three volumes per trial. In previous DecNef research, a common practice involves utilizing volume averaging to improve the signal-to-noise ratio during decoding (Taschereau-Dumouchel et al., 2018; Watanabe et al., 2017). However, as noted earlier, standard and sparse logistic regression decoders tend to yield highly bimodal or overconfident category probabilities when applied to single volumes in cross-domain classification. We reasoned that providing participants with a more moderated and continuous feedback signal might enable them to refine their strategies and learn how to self-regulate their brain activity more effectively. To illustrate, consider a scenario where a specific strategy doesn’t accurately reflect the intended pattern (i.e., the probability of decoding the target class is only marginally significant). However, if the classifier tends to provide extreme or overly confident predictions, then this strategy would appear to be just correct for the participants. In our pilot studies, we investigated the issue of standard and sparse decoders showing overly confident predictions for out-of-sample distributions. We conducted a specific analysis to evaluate the generalization from perception to imagery data used in the decoder construction. Our goal was to assess whether the average decoding probability, computed by independently decoding volumes and then averaging their probabilities, demonstrated reduced bimodality compared to the probabilities obtained using the conventional method of averaging all volumes before decoding in each trial. We elected to decode each volume and then compute the average probability as feedback signal because the analysis indicated that the probability distributions were less binomial/overconfident in this case (see Margolles et al., 2023 for a comparison of both averaging techniques).

### Visual search task

After completing the final two DecNef training runs, we conducted a visual search task while the participants were still inside the MRI scanner. The purpose was to evaluate the behavioural and neural effects of DecNef training regarding the creation of new conceptual associations. The experiment involved five fMRI runs, each with a duration of approximately 6 minutes and encompassing 56 trials per run.

Each trial started with a white fixation mark displayed on a black background for 500 ms (.37*◦*) (Figure 1C). Subsequently, participants were randomly presented with either the targeted hiragana by DecNef or a control hiragana they had not seen before for 1.5 seconds. The hiragana was presented at fixation and subtended 1.29 *◦* of visual angle. Next, participants were briefly shown two images —an example of a dog and an example of a scissor— each spanning 8.06 degrees of visual angle. These images appeared to the left and right of the fixation point and were displayed for 150 ms. The search display was followed by the presentation of a letter indicating the target image for participants to respond to: ‘P’ for dogs (from the Spanish word *’Perros’*) and ‘T’ for scissors (from the Spanish word *’Tijeras’*). The letter remained on the screen for 1750 ms, which also served as the response deadline for participants. Participants were asked to use their dominant hand (i.e., right) to indicate the location of the picture corresponding to the target category on a MRI-compatible response pad as quickly and accurately as possible. Specifically, they were instructed to press the button on the right or left side of the pad that corresponded to the location of the picture. To align the orientation of the buttons with their left and right hand positions, they were advised to hold the response pad horizontally. In addition, participants were instructed to use their index and middle fingers of their dominant hand to press the response pad buttons. To ensure that all variables were counterbalanced in the experiment, a total of 56 trials were conducted. These trials encompassed all eight potential combinations of the three variables: hiragana (targeted vs. novel), picture position (dogs left - scissors right vs. scissors right - dogs left), and Search target (dogs vs. scissors). Each combination was presented an equal number of times (7 trials each). The order of presentation for each combination was randomized across runs and participants. A jittered ITI ranging from 2.5 to 4.5 seconds with a pseudo-exponential distribution was used between trials, with 50% of ITIs set to 2.5 seconds, 25% to 3 seconds, 11% to 3.5 seconds, 7% to 4 seconds, and 7% to 4.5 seconds. This was done to help estimate BOLD responses across trials in univariate off-line analyses (Ollinger, Corbetta, & Shulman, 2001). During the ITI, a red fixation point was displayed (.37*◦*). Before the experimental session began, participants completed some practice trials without any hiragana presentation, where a fixed ITI of one second was used.

The participants in the control group performed the same visual search task outside the fMRI scanner, but we ensured that stimulus presentation and event timing were consistent with the experimental conditions of the DecNef group. Also, we matched the exposure to the images in both experimental groups prior to performing the search task. Accordingly, the control group was presented with a block-wise design where images of dogs and scissors were shown, similar to the session in the DecNef group used to build the classifier. Next, they practiced the visual search task without any hiragana presentation. Finally, they completed the visual search task using the same timings and jitters as those used by the fMRI participants. Additionally, they listened to scanner noise through headphones while carrying out the task. Previous studies assessing the processing advantage for living items (i.e. animals) over non-living stimuli indicates that low-level visual features, such as power spectrum, luminance, and contrast, cannot account for this processing advantage (Wichmann, Braun, & Gegenfurtner, 2006; Wichmann, Drewes, Rosas, & Gegenfurtner, 2010) and that higher-level semantic categorization processes are most critical (He & Cheung, 2019; Moon, He, Ditta, Cheung, & Wu, 2022). However, Thorpe, Gegenfurtner, Fabre-Thorpe, and Buèlthoff (2001) showed how the detection of animal items dropped with increasing eccentricity levels in peripheral vision. Hence we controlled for the visual angles of the search stimuli across the DecNef and control groups.

### Questionnaires

After completing the DecNef training but before the visual search task, participants responded to a series of questions related to the effort required to maximize monetary rewards and their perceived ability to manipulate the circle’s size. We used a Likert scale from 1 to 10, where the lowest value signified *minimal effort/control* and the highest value represented *significant effort/control*. We also asked them about the strategies they used during DecNef sessions and any hypotheses they had concerning the experimental objectives.

After the visual search task, we revisited the participants’ understanding of the experiment’s goals and asked them to make a guess about the category targeted by DecNef, which could be either dogs or scissors. They were also asked to indicate their level of confidence in their answer using a 4-point Likert Scale (0 - *Chance*; 1 - *Not so sure*; 2 - *Sure*; and 3 - *Really sure*). Lastly, participants were given a debriefing regarding the true objectives of the experiment.

### Off-line whole-brain searchlight analysis of DecNef training

We used searchlight analyses to examine the probability of detecting the target category throughout the DecNef sessions. The goal of this analysis was to assess the role of the bilateral fusiform gyrus during DecNef training, in addition to the involvement of other brain regions in representing the target class. Similar to previous searchlight analyses, we utilized a standard logistic regression classifier with L2 regularization to train on the perceptual data and calculate the probability of decoding the Dog class across all DecNef trials. While our custom logistic regression classifier with Binary Cross-Entropy loss would have been the most suitable option for this analysis, given that it was utilized during DecNef training in our ROI, we decided against using it for searchlight analyses. This decision was based on the substantial computational expenses and the extended training duration it would demand for each searchlight. The searchlight analysis was conducted using the BrainIAK library (Kumar et al., 2021) for each participant. The resulting searchlight maps were then registered from the functional to the participant’s T1-weighted structural space using *epi_reg* FSL tool (Jenkinson, Beckmann, Behrens, Woolrich, & Smith, 2012), and from the structural space to the Montreal Neurological Institute (MNI) standard space (MNI-152 template, 2 * 2 * 2 mm^3^) using FMRIB’s Linear Image Registration Tool (Jenkinson, Bannister, Brady, & Smith, 2002; Jenkinson et al., 2012). A non-parametric one-sample t-test was used to identify the brain regions where the average probabilities were significantly above the chance level of 0.5 across participants. This contrast was calculated using FSL Randomise with Threshold-Free Cluster Enhancement (TFCE cluster correction) and 10,000 random permutations (S. M. Smith & Nichols, 2009). To increase the statistical power of the analyses, a spatial FWHM Gaussian Kernel of 5 mm was used for variance smoothing. The Family-wise Error (FWE) correction was applied to adjust for multiple comparisons. We obtained a whole-brain corrected statistical image, which was then thresholded to display only statistically significant clusters (p < 0.05).

### Linear Mixed Effects Modelling (LMM) analyses

Mixed effects modeling was used to examine participant’s learning effects throughout the DecNef training, namely, whether participants could progressively enhance their ability to induce neural patterns related to the target category over the course of training. The design involved repeated measures for each participant and a hierarchical structure comprising 2 groups of hiraganas, 18 participants, 4 sessions, 8 runs and 18 trials. In cases of non-independence of the data, multilevel models, specifically Linear Mixed-effects Models (LMMs), have gained popularity in psychological research. LMMs offer several advantages, such as accounting for both within-subject and between-subject variability, handling missing data, and examining relationships between variables at different hierarchical levels. Traditional analysis methods, such as repeated-measures ANOVA, often aggregate data within a single condition or hierarchical level, assuming data independence, which can lead to a loss of information and statistical power. In comparison, LMMs can capture variability in the observations by leveraging all levels of data, using both fixed and random effects modeling (Meteyard & Davies, 2020; Singmann & Kellen, 2019). To perform the linear mixed model analysis, we used the *lme4* package (Bates, Mächler, Bolker, & Walker, 2014) in the R programming language (R Core Team, 2021). Further, we employed the lmerTest package (Kuznetsova, Brockhoff, Christensen, et al., 2015) to determine the statistical significance of our results. This package estimated p-values for mixed models using Satterthwaite’s method to approximate the degrees of freedom for t and F tests.

### LMM analysis of DecNef induction probabilities

Here we used the decoding probability of each trial as the dependent variable and the global trial index (1 to 468) as a fixed effect predictor. This independent variable captures the progression of performance across trials and is critical for assessing the learning effects. We also sought to account for the potential variability in the data arising from the hierarchical structure of our study. To achieve this, we introduced a random intercept for every distinct combination of the hiragana group, participant (1 - 18), session (1 - 4), and run (1 - 8) levels. Incorporating additional time-dependent information from the run or session serve as a broader contextual factor to partial out learning contributions that are trial specific from those that develop more gradually, including fatigue effects unrelated to the trial-wise learning trend. Specifically, we used the LMM formula: ‘*lmer(trial_probability ∼ global_trial_index + (1 | hiragana group / participant / session / run))*’, to conduct our analyses in R. The dependent variable was ‘*trial_probability* ‘, the independent variable was ‘*global_trial_index* ‘, and the term ‘*(1 | hiragana group / subject_nr / session / run)*’ represented the random intercepts. Importantly, the global trial index variable was treated as a continuous numeric variable, while the hiragana group, participant, session and run variables were categorical variables with discrete levels.

### LMM analysis of behavioral data from the visual search task

To examine the effects of the DecNef training on the visual search task, a second LMM was utilized to analyze the reaction times (RTs) of the correct responses. RTs outside of 3 standard deviations from each participant’s average RT were excluded. The factors hiragana (targeted vs. control), search target (dogs vs. scissors) and their interaction were set as fixed effects. Additionally, we included each participant in the DecNef group as a random effect to account for inter-subject variability with the following formula: *lmer(RTs ∼ hiragana + search target + hiragana * search target + (1 | participant))*. In this model, the control hiragana level was used as the reference level for the hiragana factor, while the scissors level was used as the reference level for the search target factor.

Moreover, in order to compare the RTs between the DecNef and control groups, we combined the RTs across the levels of the hiragana factor (targeted vs. novel), as this factor did not apply to the control group. Then, we conducted a similar LMM using the following formula: *lmer(decision RTs ∼ search target + experimental group + search target * experimental group + (1 | participant))*. In this model, the scissors level was used as the reference level for the search target factor, while the control group level was used as the reference level for the experimental group factor.

### Univariate fMRI analyses performed on the visual search task

Although the bilateral fusiform gyrus was our a priori ROI, it is possible that DecNef-related neural changes might also occur in various regions within the ventral visual pathway. Hence, our initial approach involved conducting whole-brain analyses to explore group effects during the visual search task.

The FSL suite tools were used for conducting univariate analyses (FMRIB Software Library v6.0 available at fsl.fmrib.ox.ac.uk/fsl/fslwiki/FSL) (Jenkinson et al., 2002; S. M. Smith et al., 2004). First, the T1-weighted structural image was cropped by employing the FSL Robust Field of View (FOV) method to remove the neck and lower head slices. The resulting image was then skull-stripped using the Brain Extraction Tool. Next, RAW functional data in NifTi format was preprocessed using fMRI Expert Analysis Tool (FEAT) (Woolrich, Behrens, Beckmann, Jenkinson, & Smith, 2004; Woolrich, Ripley, Brady, & Smith, 2001). To ensure signal stabilization, the first five volumes of each functional run were discarded and the MCFLIRT algorithm was applied to correct for head-motion (Jenkinson et al., 2002). Non-brain voxels were removed through functional brain extraction, and the data was smoothed with a 5 mm full-width-half maximum (FWHM) gaussian kernel. Additionally, high-pass temporal filtering was performed with sigma = 60s. Finally, the functional data was linearly registered to the high-resolution structural brain image using a Boundary Based Registration (BBR) cost function. Then, linear affine transformations with 12 parameters were used to transform the data into the standard MNI-152 space (2*2*2 mm^3^).

Following the preprocessing of the data, we carried out hierarchical general linear modeling using FMRIB’s Improved Linear Modelling (FILM). For each visual search run, the first level analysis modeled the time-series of each voxel by incorporating four regressors (Woolrich et al., 2001): targeted hiragana - dogs target, targeted hiragana - scissors target, control hiragana - dogs target, and control hiragana - scissors target. Explanatory variables in the GLM were specified based on the onset of hiragana in each trial across the different experimental conditions. These onset times were adjusted to account for the 5 heat-up volumes that were discarded in the preprocessing stage. A double-gamma canonical hemodynamic response function was used to model each trial in the GLM beginning with the onset of the hiragana. To address motion-related confounding factors, the standard and extended motion parameters estimated by MCFLIRT were included as covariates. Additionally, to account for temporal autocorrelation in the voxel time-series, prewhitening was performed on the data. To improve the model, we incorporated temporal derivatives and filtering to the design matrix, which helped to reduce the effects of lags in the onset of the H.R.F and minimize the presence of low-frequency noise over time. Four regressors were used: control hiragana - dogs target, control hiragana - scissors target, targeted hiragana - dogs target, targeted hiragana - scissors target. For each run, we calculated 4 contrasts of parameter estimates (COPES), namely, control hiragana > targeted hiragana (i.e., 1, 1, -1, -1); control hiragana < targeted hiragana (i.e., -1, -1, 1, 1); control hiragana - dogs target > targeted hiragana - dogs target & control hiragana - scissors target < targeted hiragana - scissors target (i.e., 1, -1, -1, 1); control hiragana - dogs target < targeted hiragana - dogs target & control hiragana - scissors target > targeted hiragana - scissors target (i.e., -1, 1, 1, -1). Then, we combined the first-level COPEs files into a second-level fixed-effects model. Finally, group-based, third-level analyses (Woolrich, Behrens, Beckmann, et al., 2004; Woolrich, Behrens, & Smith, 2004) were performed using the second-level COPEs files obtained in the previous step for each individual. To address inter-subject variability within each group, a mixed-effects analysis was conducted using the FMRIB Local Analysis of Mixed Effects (FLAME) 1 + 2 algorithm. Group-based analyses used cluster based thresholding, with a voxel-wise threshold of Z = 3.1 and a cluster significance threshold of P = 0.05, corrected for multiple comparisons using Gaussian Random Field Theory. The uncorrected contrast of parameter estimates and zstats for the different contrasts are available at: https://pedromargolles.github.io/pyDecNef/.

After completing the whole-brain analyses, we narrowed our focus to the bilateral fusiform gyrus as our target ROI. This was achieved by applying the GLM to binary masks of the fusiform gyrus in MNI space. These masks were generated based on the off-line searchlight analyses discussed earlier, which identified the precise areas surrounding the fusiform gyrus that demonstrated increased probabilities of the dogs category across the DecNef sessions. Additionally, we assessed the activation levels within each individual’s fusiform masks employed during DecNef training. For each participant, we calculated the mean activation within these fusiform masks across the four different conditions (control hiragana - dogs target, control hiragana - scissors target, targeted hiragana - dogs target, and targeted hiragana - scissors target). We then analyzed these averages using a repeated measures ANOVA.

## Results

First, we report the results of the searchlight analyses carried out on perception as well as on the the cross-domain generalization from perception to imagery data to draw an individual mask involving the fusiform cortex for DecNef training (see Methods). The stimulus class (dogs vs. scissors) could be reliably decoded through the entire ventral visual pathway and beyond during the perceptual scans (see Figure 2A). The cross-domain searchlight analysis revealed significant decoding clusters in the bilateral fusiform gyrus, calcarine cortex, precuneus, supramarginal gyrus, and neighboring regions (Figure 2B).

**Figure 2:**
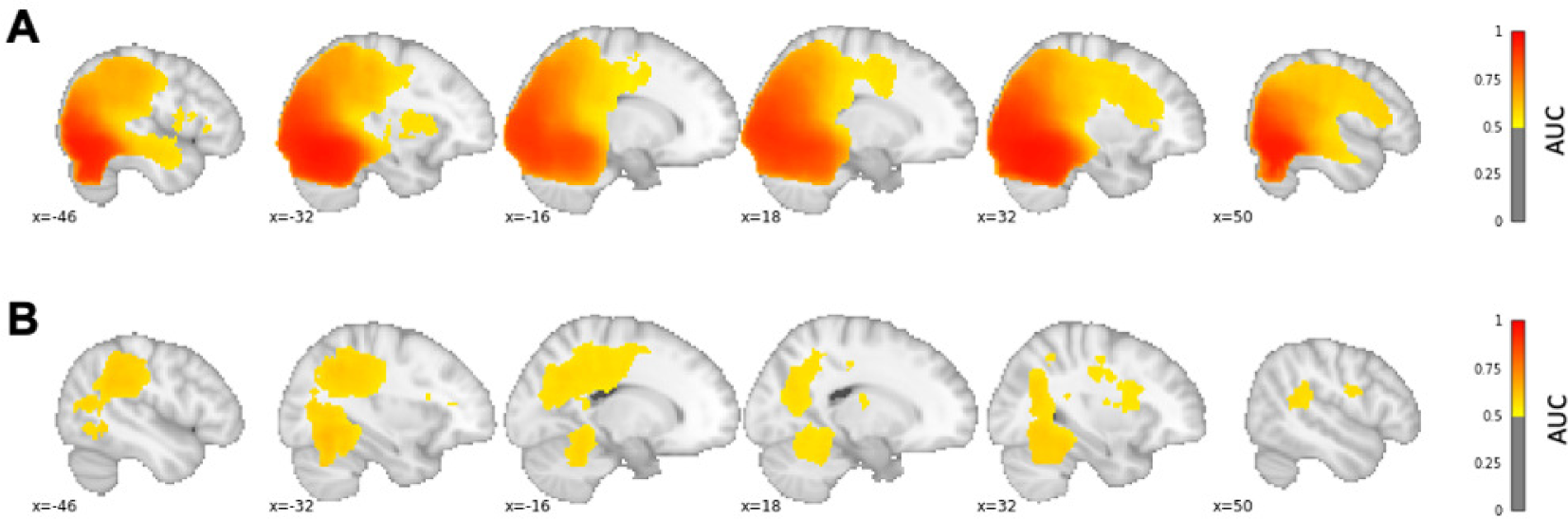
(A) Searchlight decoding results during perception. (B) Cross-domain searchlight decoding results from perception to mental imagery. The average AUC across participants is illustrated in MNI space.

Then, we assessed the participants’ ability during DecNef training to induce brain activity patterns in the target bilateral fusiform ROIs corresponding to a living world referent, which unbeknownst to them referred to the dogs pictures. A LMM analysis demonstrated significant learning effects across DecNef sessions regarding the induction of the target pattern in the fusiform cortex (Figure 3) [*β* = 7.833*e^−^*^05^, SE = 3.281*e^−^*^05^, 95% CI Lower = 1.338*e^−^*^05^, 95% CI Upper = 1.428*e^−^*^04^ (units: proportion points), p = .019]. This and subsequent analyses were repeated by excluding the three subjects who did not have a bilateral fusiform ROI, and the main experimental results were replicated (see ‘Control analyses’ section for more details).

**Figure 3:**
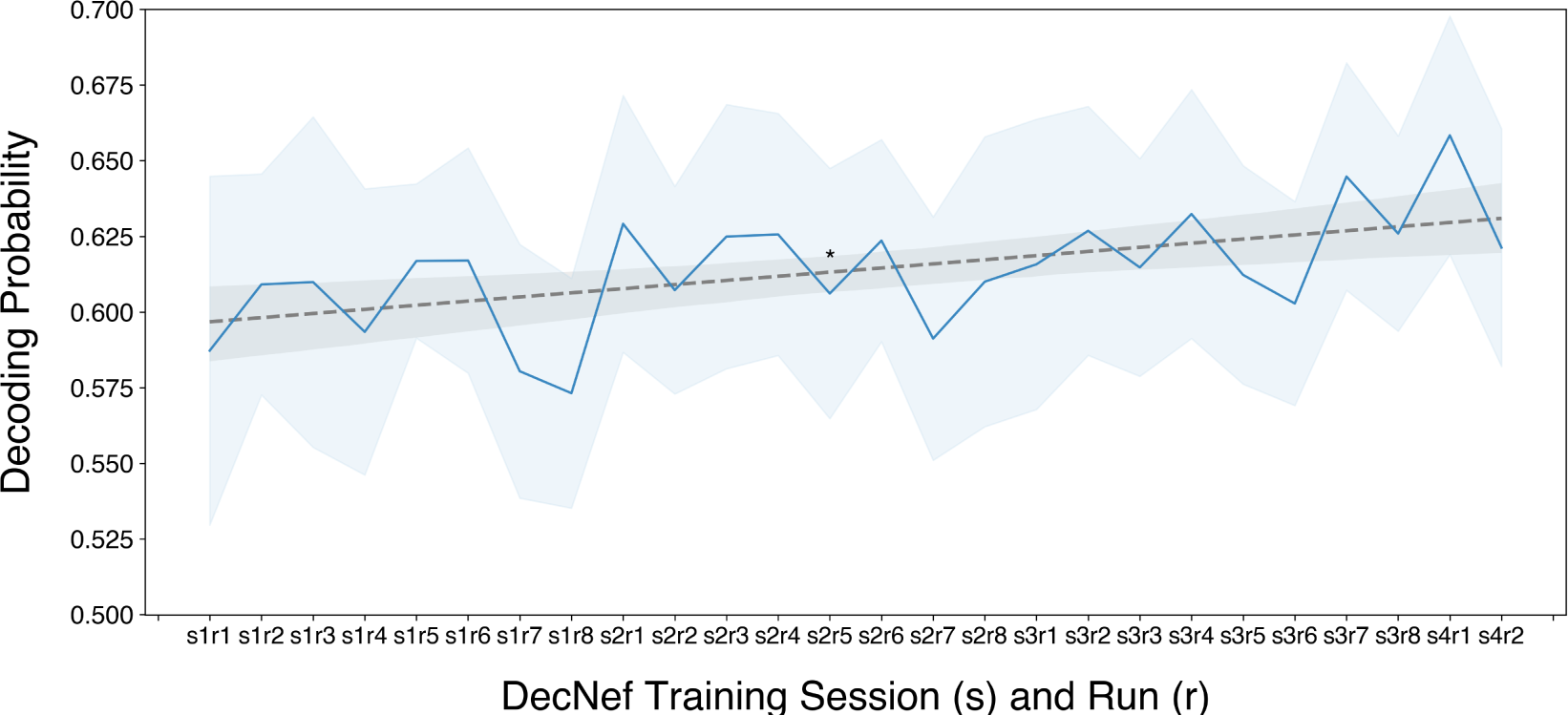
Learning effects throughout DecNef training. The figure represents the average decoding probability of dogs category in the bilateral fusiform at the run level across participants. * p = 0.019. Error bars represent 95% CIs, calculated using the Cousineau–Morey method (Cousineau et al., 2005; Cousineau & O’Brien, 2014).

We also investigated whether brain regions beyond the bilateral fusiform played a role in the induction of the relevant target category during DecNef. We performed an off-line searchlight decoding analysis using the entire brain dataset collected during the DecNef sessions. The findings revealed the involvement of additional brain regions in inducing the target pattern, including the early visual cortex (specifically, the lingual and calcarine sulcus), the precuneus, and the posterior cingulate gyrus (see Figure 4). These brain regions overlap with the areas showing better cross-domain searchlight generalization from perception to imagery during the construction of the classifier.

**Figure 4:**
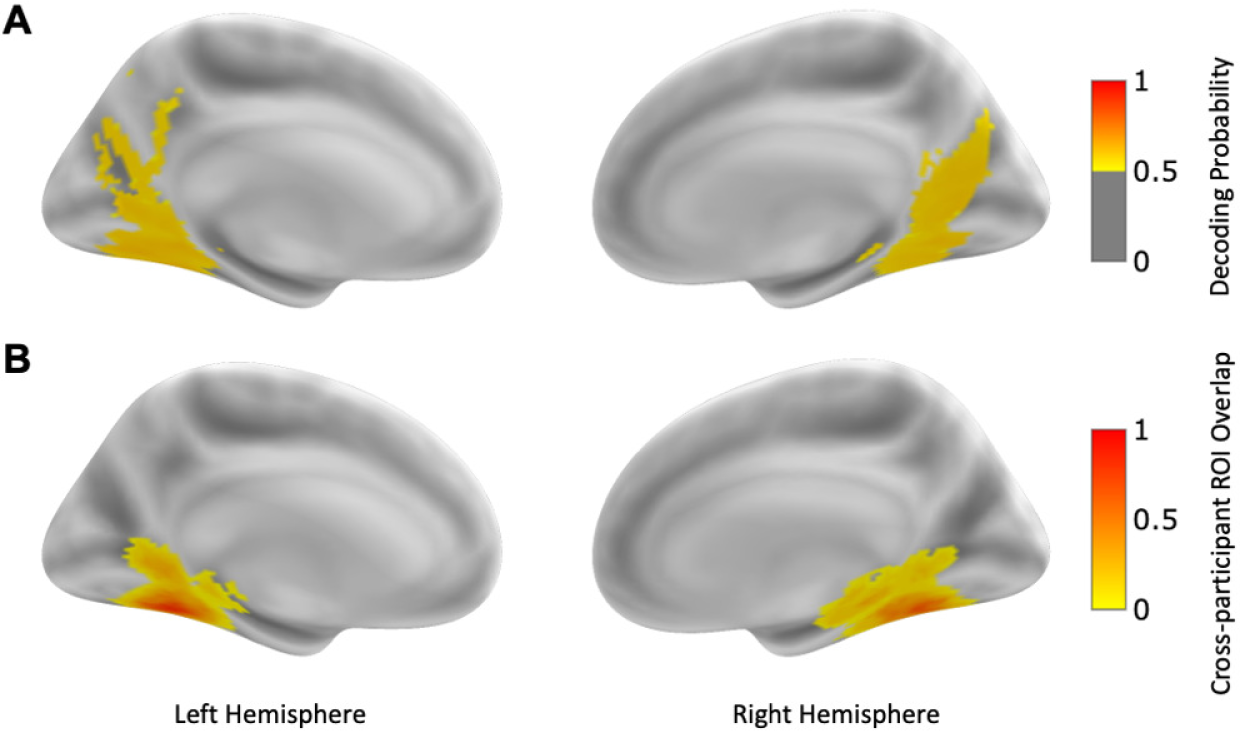
(A) Searchlight maps in MNI space where the probability of inducing the target category (i.e., dogs) across DecNef sessions exceeded the nominal 0.5 chance-level. (B) The heatmap illustrates the intersection of ROIs targeted across participants, calculated in MNI space.

### Visual search task: behavioural results

To assess the impact of DecNef training on the visual search task, we conducted a LMM model on the search RTs of the correct responses (see Methods). On average, 2.53% of trials were removed across participants due to response errors (SD = 1.9%). We also excluded outlier RTs that fell outside of 3 standard deviations from the each participant average RT (mean: 1.17% of trials; SD = 0.71%). The results showed that search RTs were significantly longer for dogs items compared to scissors [*β* = 22.053, SE = 5.650, 95% CI Lower = 10.981, 95% CI Upper = 33.126 (units: miliseconds), p < .001]. However, the main effect of the hiragana (targeted vs neutral) on search RTs was not significant [*β* = 4.795, SE = 5.647, 95% CI Lower = -6.272, 95% CI Upper = 15.863 (units: miliseconds), p = .396]. The interaction between the factors hiragana and search target was also not significant [*β* = -10.836, 95% CI Lower = -26.501, 95% CI Upper = 4.829 (units: miliseconds), p = .175], although, numerically, search for dogs was faster when cued by the targeted hiragana. Figure 5A depicts these results.

**Figure 5:**
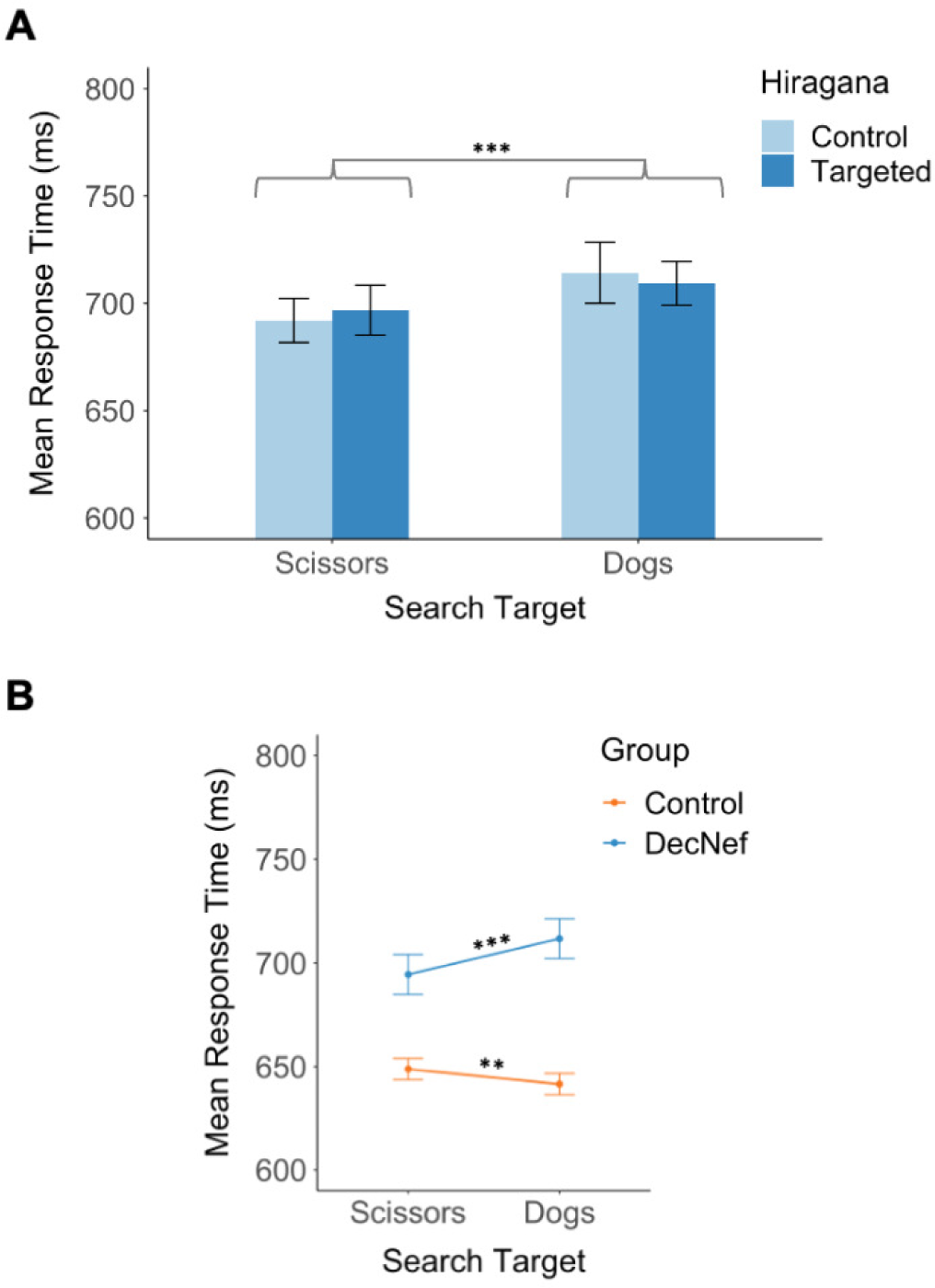
Results of the behavioral assessment in the visual search task. (A) DecNef group: visual search reaction times (RTs) as a function of the hiragana and search target factors. (B) Search RTs based on the search target across DecNef and control groups. Error bars represent 95% CIs, calculated using the Cousineau–Morey method (Cousineau et al., 2005; Cousineau & O’Brien, 2014). ** p < 0.005, *** p < 0.001.

The slower search RTs for dogs items in DecNef participants is surprising, since previous research has demonstrated the faster recognition of living categories compared to non-living counterparts (He & Cheung, 2019; Moon et al., 2022; New, Cosmides, & Tooby, 2007; Wang, Tsuchiya, New, Hurlemann, & Adolphs, 2015). This prior evidence provides a critical baseline to assess and interpret the behavioral effects of DecNef on the visual search task. In line with this, a control group (N=36) that did not undergo DecNef but performed the same task under matching conditions (see Methods), showed the opposit search bias. Namely, faster search for living (i.e., dogs) compared to non-living targets (i.e., scissors). This dissociation indicates that the effect of DecNef is selective to the target class.

Data analyses for the control group followed the same pipeline. RTs of correct search responses were analysed (Mean error = 1.82%, SD = 1.61%). Similar to the DecNef group, outlier RTs beyond 3 standard deviations from the each participant average RT were removed (Mean = 1.23%, SD = .53%). In order to compare the search RTs across DecNef and control groups, the RT data were collapsed across the levels of the hiragana factor, since this factor was not applicable to the control group. The results of the LMM analysis showed no significant differences in search RTs across the DecNef and control groups [*β* = 40.029, SE = 30.423; 95% CI Lower = -13.534, 95% CI Upper = 105.594 (units: miliseconds), p = .136]. Crucially there was a robust interaction between group and the search target factor [*β* = 24.058, SE = 4.563; 95% CI Lower = 15.115, 95% CI Upper = 33.001 (units: miliseconds), p < .001]. Simple effect analyses indicating that participants in the control group showed significantly faster RTs to pictures of dogs compared to scissors [Contrast: scissors - dogs, *β* = 7.42, SE = 2.60 (units: miliseconds), p = .004], while the DecNef group showed the opposite pattern [Contrast: scissors - dogs, *β* = -16.64, SE = 3.75 (units: miliseconds), p < .001] (Figure 5B). Additionally, we observed that the DecNef participants displayed slower search RTs for dogs compared to the control group [Contrast: control - DecNef, *β* = -70.1, SE = 30.4 (units: miliseconds), p < .021] but no differences in search RTs for scissors [Contrast: control - DecNef, *β* = -46.0, SE = 30.4 (units: miliseconds), p = .130].

We also analyzed the search error rate in the DecNef group by means of a repeated measures ANOVA in JASP (JASP Team, 2023) with hiragana and search target as factors. No significant effects of DecNef were observed on the error rate of the search responses, namely for the hiragana [F(1, 17) = .029, p = .868; *η*^2^*_p_* = .002] or the search target factor [F(1, 17) = .022, p = .883; *η*^2^*_p_* = .001]. There was no significant interaction between the two factors [F(1, 17) = .471, p = .502, *η*^2^_*p*_ = .027]. Further, no significant differences in error rate as a function the search target were found in the control group [t = -0.695, p = .492]. Finally, to compare the error rates between the two experimental groups, another ANOVA was conducted with the search target as a within-subjects factor and experimental group as a between-subjects factor. No significant effects were observed for the search target [F(1, 52) = .265, p = .609, *η*^2^_*p*_ = .005], Experimental Group [F(1, 52) = 1.972, p = .166, *η*^2^_*p*_ = .037], or the interaction between the two factors [F(1, 52) = .061, p = .805, *η*^2^*_p_* = .001].

Lastly, we run a correlation analysis between the individual mean probability of inducing the target category during DecNef training (Amano et al., 2016; Cortese et al., 2016) and the individual effect of DecNef on visual search performance (RT for Dog targets - RT for Scissor targets). We run 3 different analyses: (1) A correlation involving the mean probability across all DecNef training runs [Pearson: r = .004, p = .984; Spearman: ρ = .091, p = .717]; (2) A correlation utilizing the mean likelihood from the last two sessions [Pearson: r = -.079, p = .753; Spearman: ρ = -.077, p = .760]; (3) A correlation based on the mean likelihood from the final session [Pearson: r = -.086, p = .734; Spearman: ρ = -.007, p = .977]. All these analyses resulted in non-significant correlations.

### Visual search task: fMRI results

### Univariate analysis

Potential changes in brain activity driven by DecNef training were assessed through univariate analyses of the fMRI data acquired while participants performed the visual search task. It is well established that perceptual expectations or implicit memory templates have a strong effect on neural responses during perceptual decisions and visual search tasks (Soto et al., 2008; Soto et al., 2007; Summerfield, Trittschuh, Monti, Mesulam, & Egner, 2008). These effects have been linked to the phenomenon of neural repetition suppression, whereby neural responses decrease following the presentation of an expected item or following the presentation of an item that matches an implicit memory trace. Accordingly, based on prior studies assessing the interplay between memory and visual selection (Soto, Greene, Chaudhary, & Rotshtein, 2012; Soto et al., 2007), here we hypothesized that if the hiragana was incepted with the perceptual representations of the dogs class, then a neural repetition suppression response would be triggered by the subsequent presentation of the dogs image in the search task, relative to when a neutral (non-incepted) hiragana preceded the search display. Nonetheless, we did not observe any statistically significant clusters of activity that aligned with our hypothesis, both in terms of whole-brain analysis and within a bilateral fusiform region ROI in MNI space, as determined by the off-line searchlight analyses of DecNef training data. Further, we computed the average activity in the bilateral fusiform ROI for each participant in native space and performed a repeated measures ANOVA with the following factors: hiragana (targeted vs. control) and search target (dogs vs. scissors). There were no significant main effects of the factors associated with the hiragana [F(1, 17) = .181, p = .676, *η*^2^*_p_* = .011] or the search target [F(1, 17) = .019, p = .893, *η*^2^*_p_* = .001] and no significant interaction [F(1, 17) = .312, p = .584, *η*^2^_*p*_ = .018].

### Multivariate analysis

We further conducted a decoding analysis on the visual search fMRI data. We employed the previously trained perceptual classifiers to predict the probability of the dogs class for each individual trial in the search task. This process involved calculating the probability of the volume aligning with the corresponding hemodynamic peak during each presentation of hiragana, specifically within the temporal range of 4 to 6 seconds after the hiraganas’ onset. The fundamental premise underlying this analysis is that successful conditioning through DecNef should enhance the probability of categorizing a trial as dogs when presented with the targeted hiragana cue, as opposed to the control hiragana cue. To quantify this, we computed the average probability of dogs across runs per participant, and conducted a paired t-test to compare these probabilities between the targeted and control hiragana conditions. We observed no significant differences between the targeted hiragana (Mean = .769, SE = .011) and the control hiragana (Mean = .773 SE = .012) [t(17) = 1.082, p = .294].

### PCA analysis

So far, the results indicate that DecNef training did not result in associative knowledge between the hiragana and the perceptual content represented in the decoded multivoxel activity pattern in the fusiform. However, the behavioural results indicate that the representation of the target concept changed following DecNef training. We then further examined representational changes induced by DecNef training by means of a Principal Component Analysis (PCA) of the fusiform activity patterns across DecNef runs (Tipping & Bishop, 1999). This approach has been previously used to characterize the brain representation of both sensory (Deitch, Rubin, & Ziv, 2021; Mazor & Laurent, 2005), and motor information (Ahrens et al., 2012; Churchland, Cunningham, Kaufman, Ryu, & Shenoy, 2010) in cortex.

The analysis was performed in Scikit-learn (Pedregosa et al., 2011) on an individual basis. Specifically, Principal Components (PCs) were identified from the dogs perceptual samples used to train the decoder. Prior to performing the PCA, trial examples were standardized by subtracting the mean and dividing by the standard deviation. Subsequently, the DecNef data were projected onto the space identified by the PCs. Then, a linear Support Vector Regression (SVR) was used to test for linear trends in the representation defined by the PCs across the runs of DecNef training for each participant. The linear SVR used the PCs of the DecNef data to predict the indices of the DecNef training runs. The SVR was cross-validated using a 100-fold cross-validation procedure before deriving the average variance explained across folds. We run the analysis with 25 PCs based on prior analysis of DecNef induction dataset (Shibata et al., 2019) and also with 10 and 3 PCs in order to estimate the robustness of the results.

The results of the PCA/SVR analysis in the fusiform ROI targeted by DecNef showed significant changes in the representation of the PCs across DecNef runs. This result was consistent across different analyses performed with different numbers of PCs [25 PCs: t(17) = 2.953, p = .004; 10 PCs: t(17) = 2.34, p = .016; 3 PCs: t(17) = 1.758, p = .048].

The same PCA/SVR analysis was performed in four control ROIs. The inferior frontal and inferior parietal ROIs were selected as control regions because their activity patterns did not significantly predict the target class in the searchlight analysis Figure 4. We also included a second pair of control regions from the searchlight analysis, namely, posterior cingulate and precuneus, that showed above chance probability of decoding the target class.

We directly compared the variance explained in the fusiform and the control ROIs by means of 5 (ROI) x 3 (number of PCs) repeated measures ANOVA using Greenhouse-Geisser for the sphericity correction correction. There was a main effect of the ROI factor [F(4, 68) = 4.235, p *<* .025, *η*^2^*_p_* = .199]. There was no main effect of number of PCs [F(2, 34) = 2.522, p = .126, *η*^2^*_p_* = .129] and no interaction between the number of PCs and ROIs [F(8, 136) = 2.464, p = .105, *η*^2^*_p_* = .127]. Figure 6 depicts the pattern of results. Additional post-hoc tests, using Holm correction for multiple comparisons, showed that the variance explained in the fusiform was significantly higher compared to the other ROIs [i.e. inferior frontal: t = 3.421, p = .011; inferior parietal: t = 3.381, p = .011; posterior cingulate: t = 3.166, p = .019] except the precuneus [t = 2.054, p = .307].

**Figure 6:**
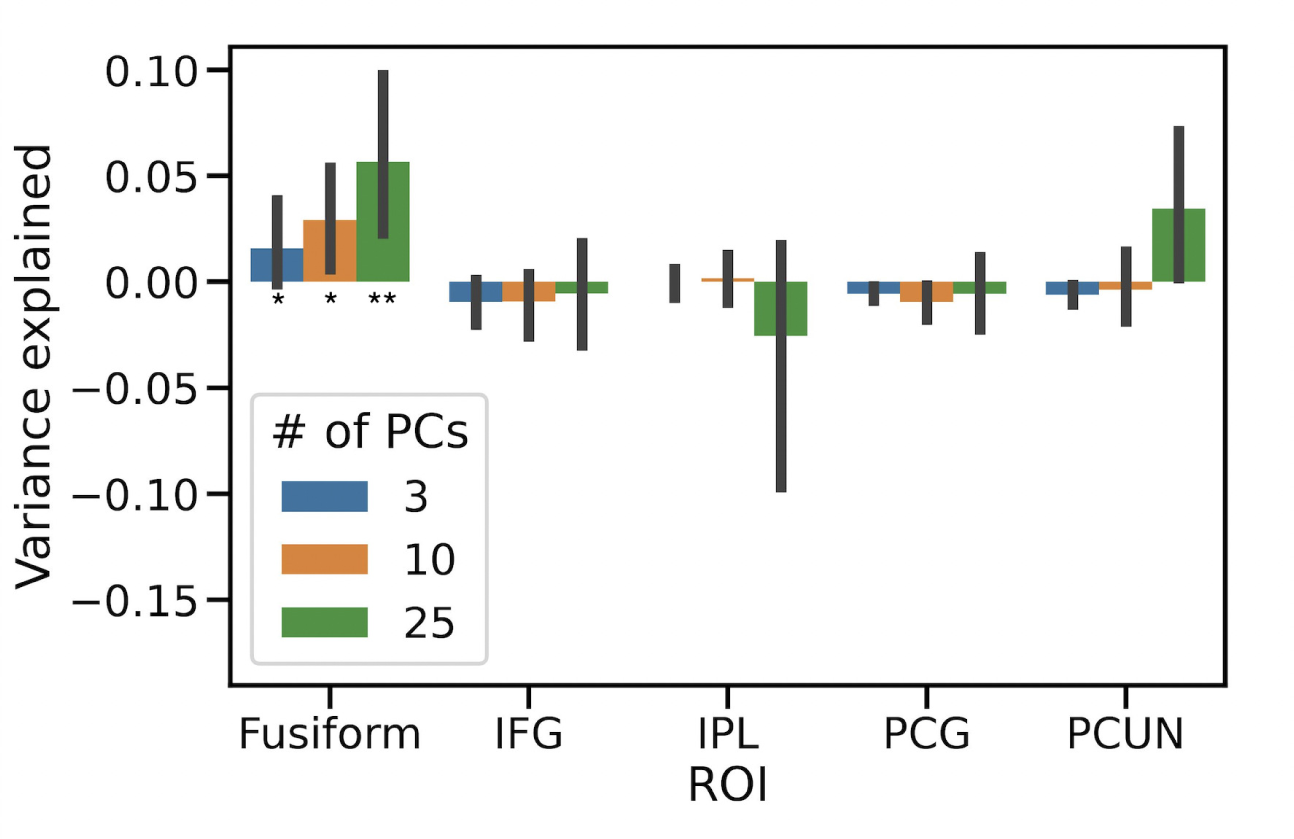
Variance explained by the linear Support Vector Regression (SVR) analysis as a function of ROI and number of Principal Components (PCs). The ROIs include: Fusiform, Inferior Frontal Gyrus (IFG), Inferior Parietal Lobe (IPL), Paracingulate Gyrus (PCG), and Precuneus (PCUN). Error bars represent 95% CIs, calculated using the Cousineau–Morey method (Cousineau et al., 2005; Cousineau & O’Brien, 2014). * p < 0.05, ** p < 0.005

### Control analyses

We repeated a new set of LMM (Linear Mixed Models) analyses after excluding the three participants who did not specifically engage the bilateral fusiform. Our analysis revealed significant learning effects across DecNef sessions in terms of inducing the target patterns in the fusiform cortex [*β* = 8.150*e^−^*^05^, SE = 3.750*e^−^*^05^, 95% CI Lower = 9.458*e^−^*^06^, 95% CI Upper = .000 (units: proportion points), p = .034]. In the visual search task, we replicated too the main experimental results. The DecNef group was significantly slower for dogs targets compared to scissors targets [*β* = 20.392, SE = 6.238; 95% CI Lower = 8.168, 95% CI Upper = 32.617 (units: miliseconds), p = .001], but not significant main effects on RTs were found for the hiragana [*β* = 1.702, SE = 6.236; 95% CI Lower = -10.519, 95% CI Upper = 13.923 (units: miliseconds), p = .785] nor interaction effects between hiragana and search target factors [*β* = -9.435, SE = 8.833; 95% CI Lower = -26.746, 95% CI Upper = 7.874 (units: miliseconds), p = .285]. Upon comparing the experimental groups, we observed that there were no statistically significant differences in RTs between the DecNef and control groups [*β* = 58.955, SE = 30.996; 95% CI Lower = -1.725, 95% CI Upper = 119.638 (units: miliseconds), p = .063]. However, we found an interaction effect of search target and group factors [*β* = 23.104, SE = 4.840; 95% CI Lower = 13.617, 95% CI Upper = 32.591 (units: miliseconds), p *<* .001]. Analysis of the simple effects revealed that control group participants respond faster to images of dogs than to scissors [Contrast: scissors - dogs, *β* = 7.42, SE = 2.61 (units: miliseconds), p = .004]. In contrast, those in the DecNef group exhibited the reverse trend [Contrast: scissors - dogs, *β* = -15.69, SE = 4.08 (units: miliseconds), p *<* .001]. Moreover, the DecNef participants had slower RTs for dogs when compared to the control group [Contrast: control - DecNef, *β* = -82.1, SE = 31 (units: miliseconds), p = .008]. However, there was no significant difference in RTs for scissors between the two groups [Contrast: control - DecNef, *β* = -59.0, SE = 31 (units: miliseconds), p = .057]. There were no significant differences in search accuracy between the DecNef and control groups. The univariate and multivariate fMRI analyses also revealed no significant activity differences as a function of the different trial types with the targeted vs neutral hiragana.

### Participants’ subjective reports

Participants were queried about the techniques they employed to manipulate the size of the feedback disk during DecNef training. Their responses included various strategies like mental humming, mathematical calculations, visualizing different scenarios, evoking emotions, and more. Many participants indicated that they needed to adapt and employ a range of strategies to maintain their performance over time. They expressed moderate to high levels of effort in enlarging the feedback disk, but the majority believed they had gained a high degree of control throughout DecNef training. Moreover, participants reports indicate that they were unaware of the specific research goals. Out of 18 participants, twelve participants guessed that the target category was dogs, but a Chi-squared test showed the observed frequencies were consistent with a uniform distribution and therefore unreliable identification of the target class [χ2(1) = 2, p = .157]. Confidence reports indicated low confidence levels in their guesses, even among those who guessed correctly. Tables with details regarding the individual strategies and participants beliefs about experimental goals are available at: https://pedromargolles.github.io/pyDecNef/.

## Discussion

This study aimed to covertly manipulate conceptual representations using fMRI-DecNef, thereby introducing a novel perceptual meaning into a neutral hiragana without participants’ awareness. Participants progressively improved in their ability to induce neural patterns that were similar to the target concept (i.e., dogs). This observation is noteworthy because previous DecNef studies that used a similar amount of training sessions as here (Amano et al., 2016; Cortese et al., 2016; Knotts et al., 2019; Koizumi et al., 2016; Taschereau-Dumouchel et al., 2018), did not reliably observe significant learning effects, despite observing DecNef effects on behavioral performance. Here we used an improved decoder to provide a better calibrated and more fine-grained feedback to participants (Margolles et al., 2023). This improvement contrasts with the logistic regression classifiers used in previous DecNef studies, which tended to produce overly confident and highly binomial predictions as feedback (Bai et al., 2021). These improvements in the classifier could enhance participants’ capacity to identify subtle neural changes through feedback, potentially resulting in improved self-regulation during DecNef training. These improvements represent a step forward towards refining the precision of DecNef. However, there were inter-individual differences in learning during DecNef, as commonly observed in neurofeedback studies, due to differences in attention, motivation (Kadosh & Staunton, 2019) or metacognitive abilities (Lubianiker, Paret, Dayan, & Hendler, 2022; Muñoz-Moldes & Cleeremans, 2020; Stirner et al., 2022). Future DecNef studies would benefit from developing personalised training protocols to enhance learning within each individual.

Despite the consistent learning effect detected during the DecNef training, which resulted in strong non-conscious neural representations of the target perceptual content, the behavioral results do not support the view that the hiragana was associated with the target concept. Search performance for the target category (i.e., dogs) was similar when search was primed by the hiragana targeted by DecNef, or by a control hiragana. Previous studies provided robust evidence that memory effects on visual search performance can occur when the number of items in the search display is low, namely, even with two items (Pan, Cheng, & Luo, 2012; Soto et al., 2012; Soto et al., 2007; Soto, Wriglesworth, Bahrami-Balani, & Humphreys, 2010). Thereby, it is unlikely that the absence of effects of the incepted hiragana were due to the small set size of the search display. Further, both univariate and multivariate fMRI analyses in the search task did not show any neural modulation driven by the matching between the targeted hiragana and the target category. While some prior studies indicated that associative learning can operate on visually masked and subliminal representations (Pessiglione et al., 2008; Raio et al., 2012; Seitz et al., 2009), mounting evidence has indicated otherwise (Field, 2000; Mertens & Engelhard, 2020; Shanks & John, 1994; Skora et al., 2021). Prior research supporting unconscious associative learning faced methodological criticisms, particularly regarding experiment demand characteristics and limitations in assessing participant awareness (Field, 2000; Shanks & Dickinson, 1990). Our findings indicate that while robust neural representations of the target perceptual content are induced, forming associative knowledge between these patterns and a neutral stimulus likely requires conscious access to the represented content. However, this does not necessarily exclude that others forms of associative learning, namely, involving emotional content and fear conditioning (LeDoux & Pine, 2016) may occur outside participants’ awareness (Lovibond & Shanks, 2002).

A previous study found evidence of orientation-color associations in early visual cortex induced by DecNef (Amano et al., 2016). However, our current study investigated higher-order associative learning involving an abstract symbol and a perceptual class, compared to the lower-order associations observed in the previous study. More recently, Collin and colleagues (Collin, van den Broek, van Mourik, Desain, & Doeller, 2022) showed that contextual representations of faces or houses could be induced through DecNef in the visual cortex, while participants memorized a different set of objects. The implicit neural context induced by DecNef (i.e., face or house) facilitated associative memories between the study items and the specific encoding contexts. However, in Collin and colleagues’ study, participants were explicitly instructed to think about either faces or houses during DecNef training, thereby likely implicating awareness of the association between the feedback and the mental context (i.e., whether the feedback related to faces or houses).

Notably, DecNef training significantly impacted the representation of the target category. In contrast to the control group, the DecNef group exhibited impaired selection of the target category during the search. While the control group showed a visual search advantage for living (i.e., dogs) over non-living targets (i.e., scissors), consistent with prior research (e.g., He and Cheung, 2019; Moon et al., 2022; New et al., 2007; Wang et al., 2015), the DecNef group displayed the opposite pattern of results. Specifically, they showed slower search performance for dogs compared to scissors. This dissociation suggests that DecNef’s influence is specific to the targeted category. We propose that DecNef training can lead to a reconfiguration of the neural template associated with the target category, which in turn affects search behavior. Importantly, previous studies demonstrated that visual imagery, working memory and longer-term memory contents have positive influences in the guidance of attention, enhancing selection and search performance for visual targets that match the contents held in memory (Nobre & Stokes, 2019; Reinhart, McClenahan, & Woodman, 2015; Soto et al., 2008; Tartaglia, Bamert, Mast, & Herzog, 2009). For instance, it has been shown that explicit imagery training of a visual feature facilitates its subsequent selection in a visual search task (Nobre & Stokes, 2019; Reinhart et al., 2015; Tartaglia et al., 2009). Therefore, the impairment in search performance observed for the target representation of DecNef (i.e., dogs) is not predicted by an account based on participants consciously rehearsing the target representations during DecNef.

One possible explanation is that DecNef induced representational drift of the target concept, causing a shift in alignment of neural activity patterns within a high-dimensional space over time (Rule et al., 2019). DecNef protocols require participants to explore a broad search space to find the target representation by means of diverse cognitive strategies (Watanabe et al., 2017). These strategies are conceptually distinct from the target and are also accompanied by various sources of interference, such as the presence of the hiragana in this particular case. Hence, the brain is confronted with a many-to-one mapping problem often encountered in machine learning, where a unique target can potentially be mapped from multiple patterns in multidimensional space. The repeated induction of noisy neural patterns that partially overlap with the neural representation of the target class, can consequently result in the corruption of the target representation through Hebbian learning and operant conditioning (McClelland, Thomas, McCandliss, & Fiez, 1999). The PCA analyses demonstrated that the fusiform activity patterns underwent changes in representation throughout the DecNef training runs compared with several control regions. This result provides further evidence that effects of DecNef training were more pronounced and localized within the specific targeted region, in keeping with a targeted neural plasticity model of DecNef effects (Shibata et al., 2019).

Representational drift has been observed in the hippocampus (Ziv et al., 2013) and also in cortical regions (Driscoll, Pettit, Minderer, Chettih, & Harvey, 2017), including sensory cortex (Aitken, Garrett, Olsen, & Mihalas, 2022; Deitch et al., 2021), when observers are exposed to similar stimuli across sessions and days. Representational drift is higher for more complex representations associated with naturalistic stimuli relative to simple items (i.e. oriented gratings) (Marks & Goard, 2021). Importantly, representational drift can occur despite sustained decoding accuracy of the target content from neural responses by a linear classifier (Aitken et al., 2022), as we observed in the present study.

Previous theoretical work proposed that representational drift may afford computational benefits, namely, for continual learning (Driscoll, Duncker, & Harvey, 2022). However, the causal effects of representational drift on behaviour remain to be elucidated (Sadeh & Clopath, 2022). The current findings suggest that representational drift could potentially result in undesired behavioral outcomes. Specifically, representational drift may lead to a mismatch between internal representations held in memory and the perceptual input from the external environment, thereby influencing visual selection and search behaviour.

In summary, this study elaborates on the impact of conscious awareness on different types of learning. It indicates that higher-order associative learning is more likely to depend on conscious awareness, whereas lower-level forms of re-learning, modification, or plasticity in established representations can occur without awareness, potentially through a mechanism involving neural representational drift. Additionally, this research offers novel insights for the advancement of DecNef protocols, specifically aimed at manipulating conceptual representations. These findings pave the way for further research into the potential of DecNef in personalized interventions, where the guidance provided can be used to target and improve maladaptive mental representations.

## Open Science Statement

The scripts and documentation corresponding to the Python library used for decoded neurofeedback (PyDecNef) can be found at: https://pedromargolles.github.io/pyDecNef/ (Margolles et al., 2023). The experimental data will be uploaded to OpenNeuro upon acceptance for publication.

## Acknowledgements

P.M acknowledges support from an FPI grant from the Spanish Ministry of Economy and Competitiveness. P.E. acknowledges support from the Basque Government PREDOC grant. D.S. acknowledges support from the Basque Government through the BERC 2022-2025 program, from the Spanish Ministry of Economy and Competitiveness, through the ‘Severo Ochoa’ Programme for Centres/Units of Excellence (CEX2020-001010-S) and also from project grants PID2019-105494GB-I00. The funders had no role in study design, data collection and analysis, decision to publish or preparation of the manuscript.

## Competing interests statement

The authors declare no competing interests.

## Notes

https://github.com/pedromargolles/pyDecNef

### Competing Interest Statement

The authors have declared no competing interest.

### Summary of Updates

added further discussion and control analyses

## References

Abadi, M., Agarwal, A., Barham, P., Brevdo, E., Chen, Z., Citro, C., . . . Devin, M., et al. (2016). Tensorflow: Large-scale machine learning on heterogeneous distributed systems. arXiv preprint arXiv:1603.04467.

Ahrens, M. B., Li, J. M., Orger, M. B., Robson, D. N., Schier, A. F., Engert, F., & Portugues, R. (2012). Brain-wide neuronal dynamics during motor adaptation in zebrafish. Nature, 485 (7399), 471–477.

Aitken, K., Garrett, M., Olsen, S., & Mihalas, S. (2022). The geometry of representational drift in natural and artificial neural networks. PLOS Computational Biology, 18 (11), e1010716.

Amano, K., Shibata, K., Kawato, M., Sasaki, Y., & Watanabe, T. (2016). Learning to associate orientation with color in early visual areas by associative decoded fmri neurofeedback. Current Biology, 26 (14), 1861–1866.

Bai, Y., Mei, S., Wang, H., & Xiong, C. (2021). Don’t just blame over-parametrization for over-confidence: Theoretical analysis of calibration in binary classification. In International conference on machine learning (pp. 566–576). PMLR.

Bates, D., Mächler, M., Bolker, B., & Walker, S. (2014). Fitting linear mixed-effects models using lme4. arXiv preprint arXiv:1406.5823.

Blewitt, P. (1982). Word meaning acquisition in young children: A review of theory and research. Advances in child development and behavior, 17, 139–195.

Chollet, F. (2015). Keras.[online] available at: Https://github.com/fchollet/keras. Accessed, 12 (01), 2021.

Churchland, M. M., Cunningham, J. P., Kaufman, M. T., Ryu, S. I., & Shenoy, K. V. (2010). Cortical preparatory activity: Representation of movement or first cog in a dynamical machine? Neuron, 68 (3), 387–400.

Collin, S. H., van den Broek, P. L., van Mourik, T., Desain, P., & Doeller, C. F. (2022). Inducing a mental context for associative memory formation with real-time fmri neurofeedback. Scientific Reports, 12 (1), 21226.

Connolly, A. C., Guntupalli, J. S., Gors, J., Hanke, M., Halchenko, Y. O., Wu, Y.-C., . . . Haxby, J. V. (2012). The representation of biological classes in the human brain. Journal of Neuroscience, 32 (8), 2608–2618.

Cortese, A., Amano, K., Koizumi, A., Kawato, M., & Lau, H. (2016). Multivoxel neurofeedback selectively modulates confidence without changing perceptual performance. Nature communications, 7 (1), 13669.

Cortese, A., Amano, K., Koizumi, A., Lau, H., & Kawato, M. (2017). Decoded fmri neurofeedback can induce bidirectional confidence changes within single participants. NeuroImage, 149, 323–337.

Cousineau, D. et al. (2005). Confidence intervals in within-subject designs: A simpler solution to loftus and masson’s method. Tutorials in quantitative methods for psychology, 1 (1), 42–45.

Cousineau, D., & O’Brien, F. (2014). Error bars in within-subject designs: A comment on baguley (2012). Behavior Research Methods, 46, 1149–1151.

Cox, R. (1996). Afni: Programvare for analyse og visualisering av funksjonell magnetisk resonans neuroimages. Comput Biomed Res, 29, 162–173.

Deitch, D., Rubin, A., & Ziv, Y. (2021). Representational drift in the mouse visual cortex. Current biology, 31 (19), 4327–4339.

Dijkstra, N., van Gaal, S., Geerligs, L., Bosch, S. E., & van Gerven, M. A. (2021). No evidence for neural overlap between unconsciously processed and imagined stimuli. Eneuro, 8 (5).

Downing, P. E., Chan, A.-Y., Peelen, M., Dodds, C., & Kanwisher, N. (2006). Domain specificity in visual cortex. Cerebral cortex, 16 (10), 1453–1461.

Driscoll, L. N., Duncker, L., & Harvey, C. D. (2022). Representational drift: Emerging theories for continual learning and experimental future directions. Current Opinion in Neurobiology, 76, 102609.

Driscoll, L. N., Pettit, N. L., Minderer, M., Chettih, S. N., & Harvey, C. D. (2017). Dynamic reorganization of neuronal activity patterns in parietal cortex. Cell, 170 (5), 986–999.

Field, A. P. (2000). I like it, but i’m not sure why: Can evaluative conditioning occur without conscious awareness? Consciousness and cognition, 9 (1), 13–36.

Gorgolewski, K., Burns, C. D., Madison, C., Clark, D., Halchenko, Y. O., Waskom, M. L., & Ghosh, S. S. (2011). Nipype: A flexible, lightweight and extensible neuroimaging data processing framework in python. Frontiers in neuroinformatics, 13.

He, C., & Cheung, O. S. (2019). Category selectivity for animals and man-made objects: Beyond low-and mid-level visual features. Journal of vision, 19 (12), 22–22.

Hebart, M. N., Dickter, A. H., Kidder, A., Kwok, W. Y., Corriveau, A., Van Wicklin, C., & Baker, C. I. (2019). Things: A database of 1,854 object concepts and more than 26,000 naturalistic object images. PloS one, 14 (10), e0223792.

Hofmann, W., De Houwer, J., Perugini, M., Baeyens, F., & Crombez, G. (2010). Evaluative conditioning in humans: A meta-analysis. Psychological bulletin, 136 (3), 390.

JASP Team. (2023). JASP (Version 0.17.1)[Computer software]. Retrieved from https://jasp-stats.org/

Jenkinson, M., Bannister, P., Brady, M., & Smith, S. (2002). Improved optimization for the robust and accurate linear registration and motion correction of brain images. Neuroimage, 17 (2), 825–841.

Jenkinson, M., Beckmann, C. F., Behrens, T. E., Woolrich, M. W., & Smith, S. M. (2012). Fsl. Neuroimage, 62 (2), 782–790.

Kadosh, K. C., & Staunton, G. (2019). A systematic review of the psychological factors that influence neurofeedback learning outcomes. Neuroimage, 185, 545–555.

Knotts, J., Cortese, A., Tascherau-Dumouchel, V., Kawato, M., & Lau, H. (2019). Multivoxel patterns for perceptual confidence are associated with false color detection. bioRxiv, 735084.

Koizumi, A., Amano, K., Cortese, A., Shibata, K., Yoshida, W., Seymour, B., . . . Lau, H. (2016). Fear reduction without fear through reinforcement of neural activity that bypasses conscious exposure. Nature human behaviour, 1 (1), 0006.

Kumar, M., Anderson, M. J., Antony, J. W., Baldassano, C., Brooks, P. P., Cai, M. B., . . . Huberdeau, D., et al. (2021). Brainiak: The brain imaging analysis kit. Aperture neuro, 1 (4).

Kuznetsova, A., Brockhoff, P. B., Christensen, R. H. B., et al. (2015). Package ‘lmertest’. R package version, 2 (0), 734.

Lau, H. (2022). In consciousness we trust: The cognitive neuroscience of subjective experience. Oxford University Press.

LeDoux, J. E., & Pine, D. S. (2016). Using neuroscience to help understand fear and anxiety: A two-system framework. American journal of psychiatry.

Lee, S.-H., Kravitz, D. J., & Baker, C. I. (2012). Disentangling visual imagery and perception of real-world objects. Neuroimage, 59 (4), 4064–4073.

Li, X., Morgan, P. S., Ashburner, J., Smith, J., & Rorden, C. (2016). The first step for neuroimaging data analysis: Dicom to nifti conversion. Journal of neuroscience methods, 264, 47–56.

Lovibond, P. F., & Shanks, D. R. (2002). The role of awareness in pavlovian conditioning: Empirical evidence and theoretical implications. Journal of Experimental Psychology: Animal Behavior Processes, 28 (1), 3.

Lubianiker, N., Paret, C., Dayan, P., & Hendler, T. (2022). Neurofeedback through the lens of reinforcement learning. Trends in Neurosciences.

Margolles, P., Mei, N., Elosegi, P., & Soto, D. (2023). Pydecnef: An open-source framework for fmri-based decoded neurofeedback. bioRxiv. doi:10.1101/2023.10.02.560503

Marks, T. D., & Goard, M. J. (2021). Stimulus-dependent representational drift in primary visual cortex. Nature communications, 12 (1), 5169.

Mathôt, S., & March, J. (2022). Conducting linguistic experiments online with opensesame and osweb. Language Learning, 72 (4), 1017–1048.

Mathôt, S., Schreij, D., & Theeuwes, J. (2012). Opensesame: An open-source, graphical experiment builder for the social sciences. Behavior research methods, 44, 314–324.

Mazor, O., & Laurent, G. (2005). Transient dynamics versus fixed points in odor representations by locust antennal lobe projection neurons. Neuron, 48 (4), 661–673.

McClelland, J. L., Thomas, A. G., McCandliss, B. D., & Fiez, J. A. (1999). Understanding failures of learning: Hebbian learning, competition for representational space, and some preliminary experimental data. Progress in brain research, 121, 75–80.

Mertens, G., & Engelhard, I. M. (2020). A systematic review and meta-analysis of the evidence for unaware fear conditioning. Neuroscience & Biobehavioral Reviews, 108, 254–268.

Meteyard, L., & Davies, R. A. (2020). Best practice guidance for linear mixed-effects models in psychological science. Journal of Memory and Language, 112, 104092.

Moon, A., He, C., Ditta, A. S., Cheung, O. S., & Wu, R. (2022). Rapid category selectivity for animals versus man-made objects: An n2pc study. International Journal of Psychophysiology, 171, 20–28.

Muñoz-Moldes, S., & Cleeremans, A. (2020). Delineating implicit and explicit processes in neurofeedback learning. Neuroscience & Biobehavioral Reviews, 118, 681–688.

New, J., Cosmides, L., & Tooby, J. (2007). Category-specific attention for animals reflects ancestral priorities, not expertise. Proceedings of the National Academy of Sciences, 104 (42), 16598–16603.

Nobre, A. C., & Stokes, M. G. (2019). Premembering experience: A hierarchy of time-scales for proactive attention. Neuron, 104 (1), 132–146.

Oakes, T. R., Johnstone, T., Walsh, K. O., Greischar, L. L., Alexander, A. L., Fox, A. S., & Davidson, R. J. (2005). Comparison of fmri motion correction software tools. Neuroimage, 28 (3), 529–543.

Oblak, E. F., Sulzer, J. S., & Lewis-Peacock, J. A. (2019). A simulation-based approach to improve decoded neurofeedback performance. NeuroImage, 195, 300–310.

Ollinger, J., Corbetta, M., & Shulman, G. (2001). Separating processes within a trial in event-related functional mri: Ii. analysis. Neuroimage, 13 (1), 218–229.

Page, M. M. (1974). Demand characteristics and the classical conditioning of attitudes experiment. Journal of Personality and Social Psychology, 30 (4), 468.

Pan, Y., Cheng, Q.-P., & Luo, Q.-Y. (2012). Working memory can enhance unconscious visual perception. Psychonomic Bulletin & Review, 19, 477–482.

Pedregosa, F., Varoquaux, G., Gramfort, A., Michel, V., Thirion, B., Grisel, O., . . . Duchesnay, E. (2011). Scikit-learn: Machine learning in Python. Journal of Machine Learning Research, 12, 2825–2830.

Pessiglione, M., Petrovic, P., Daunizeau, J., Palminteri, S., Dolan, R. J., & Frith, C. D. (2008). Subliminal instrumental conditioning demonstrated in the human brain. Neuron, 59 (4), 561–567.

Pitcher, D., Charles, L., Devlin, J. T., Walsh, V., & Duchaine, B. (2009). Triple dissociation of faces, bodies, and objects in extrastriate cortex. Current Biology, 19 (4), 319–324.

Raio, C. M., Carmel, D., Carrasco, M., & Phelps, E. A. (2012). Nonconscious fear is quickly acquired but swiftly forgotten. Current Biology, 22 (12), R477–R479.

Reddy, L., Tsuchiya, N., & Serre, T. (2010). Reading the mind’s eye: Decoding category information during mental imagery. Neuroimage, 50 (2), 818–825.

Reinhart, R. M., McClenahan, L. J., & Woodman, G. F. (2015). Visualizing trumps vision in training attention. Psychological science, 26 (7), 1114–1122.

Sadeh, S., & Clopath, C. (2022). Contribution of behavioural variability to representational drift. Elife, 11, e77907.

Schwarzlose, R. F., Baker, C. I., & Kanwisher, N. (2005). Separate face and body selectivity on the fusiform gyrus. Journal of Neuroscience, 25 (47), 11055–11059.

Seitz, A. R., Kim, D., & Watanabe, T. (2009). Rewards evoke learning of unconsciously processed visual stimuli in adult humans. Neuron, 61 (5), 700–707.

Sha, L., Haxby, J. V., Abdi, H., Guntupalli, J. S., Oosterhof, N. N., Halchenko, Y. O., & Connolly, A. C. (2015). The animacy continuum in the human ventral vision pathway. Journal of cognitive neuroscience, 27 (4), 665–678.

Shanks, D. R., & Dickinson, A. (1990). Contingency awareness in evaluative conditioning: A comment on baeyens, eelen, and van den bergh. Cognition and Emotion, 4 (1), 19–30.

Shanks, D. R., & John, M. F. S. (1994). Characteristics of dissociable human learning systems. Behavioral and brain sciences, 17 (3), 367–395.

Shibata, K., Lisi, G., Cortese, A., Watanabe, T., Sasaki, Y., & Kawato, M. (2019). Toward a comprehensive understanding of the neural mechanisms of decoded neurofeedback. NeuroImage, 188, 539–556.

Shibata, K., Watanabe, T., Kawato, M., & Sasaki, Y. (2016). Differential activation patterns in the same brain region led to opposite emotional states. PLoS biology, 14 (9), e1002546.

Singmann, H., & Kellen, D. (2019). An introduction to mixed models for experimental psychology. In New methods in cognitive psychology (pp. 4–31). Routledge.

Skora, L. I., Yeomans, M. R., Crombag, H. S., & Scott, R. B. (2021). Evidence that instrumental conditioning requires conscious awareness in humans. Cognition, 208, 104546.

Smith, M. D. (1978). The acquisition of word meaning: An introduction. Child Development, 950–952.

Smith, S. M., Jenkinson, M., Woolrich, M. W., Beckmann, C. F., Behrens, T. E., Johansen-Berg, H., . . . Flitney, D. E., et al. (2004). Advances in functional and structural mr image analysis and implementation as fsl. Neuroimage, 23, S208–S219.

Smith, S. M., & Nichols, T. E. (2009). Threshold-free cluster enhancement: Addressing problems of smoothing, threshold dependence and localisation in cluster inference. Neuroimage, 44 (1), 83–98.

Sorger, B., Scharnowski, F., Linden, D. E., Hampson, M., & Young, K. D. (2019). Control freaks: Towards optimal selection of control conditions for fmri neurofeedback studies. Neuroimage, 186, 256–265.

Soto, D., Greene, C. M., Chaudhary, A., & Rotshtein, P. (2012). Competition in working memory reduces frontal guidance of visual selection. Cerebral cortex, 22 (5), 1159–1169.

Soto, D., Hodsoll, J., Rotshtein, P., & Humphreys, G. W. (2008). Automatic guidance of attention from working memory. Trends in cognitive sciences, 12 (9), 342–348.

Soto, D., Humphreys, G. W., & Rotshtein, P. (2007). Dissociating the neural mechanisms of memory-based guidance of visual selection. Proceedings of the National Academy of Sciences, 104 (43), 17186–17191.

Soto, D., Wriglesworth, A., Bahrami-Balani, A., & Humphreys, G. W. (2010). Working memory enhances visual perception: Evidence from signal detection analysis. Journal of Experimental Psychology: Learning, Memory, and Cognition, 36 (2), 441.

Staats, C. K., & Staats, A. W. (1957). Meaning established by classical conditioning. Journal of experimental psychology, 54 (1), 74.

Stahl, C., Haaf, J., & Corneille, O. (2016). Subliminal evaluative conditioning? above-chance cs identification may be necessary and insufficient for attitude learning. Journal of Experimental Psychology: General, 145 (9), 1107.

Stirner, M., Gurevitch, G., Lubianiker, N., Hendler, T., Schmahl, C., & Paret, C. (2022). An investigation of awareness and metacognition in neurofeedback with the amygdala electrical fingerprint. Consciousness and Cognition, 98, 103264.

Stokes, M., Thompson, R., Cusack, R., & Duncan, J. (2009). Top-down activation of shape-specific population codes in visual cortex during mental imagery. Journal of Neuroscience, 29 (5), 1565–1572.

Summerfield, C., Trittschuh, E. H., Monti, J. M., Mesulam, M.-M., & Egner, T. (2008). Neural repetition suppression reflects fulfilled perceptual expectations. Nature neuroscience, 11 (9), 1004–1006.

Tartaglia, E. M., Bamert, L., Mast, F. W., & Herzog, M. H. (2009). Human perceptual learning by mental imagery. Current Biology, 19 (24), 2081–2085.

Taschereau-Dumouchel, V., Cortese, A., Chiba, T., Knotts, J., Kawato, M., & Lau, H. (2018). Towards an unconscious neural reinforcement intervention for common fears. Proceedings of the National Academy of Sciences, 115 (13), 3470–3475.

Thorpe, S. J., Gegenfurtner, K. R., Fabre-Thorpe, M., & BuÈlthoff, H. H. (2001). Detection of animals in natural images using far peripheral vision. European Journal of Neuroscience, 14 (5), 869–876.

Tipping, M. E., & Bishop, C. M. (1999). Probabilistic principal component analysis. Journal of the Royal Statistical Society: Series B (Statistical Methodology), 61 (3), 611–622.

Tryon, W. W., & Cicero, S. D. (1989). Classical conditioning of meaning—i. a replication and higher-order extension. Journal of behavior therapy and experimental psychiatry, 20 (2), 137–142.

Wang, S., Tsuchiya, N., New, J., Hurlemann, R., & Adolphs, R. (2015). Preferential attention to animals and people is independent of the amygdala. Social Cognitive and Affective Neuroscience, 10 (3), 371–380.

Watanabe, T., Sasaki, Y., Shibata, K., & Kawato, M. (2017). Advances in fmri real-time neurofeedback. Trends in cognitive sciences, 21 (12), 997–1010.

Wichmann, F. A., Braun, D. I., & Gegenfurtner, K. R. (2006). Phase noise and the classification of natural images. Vision research, 46 (8-9), 1520–1529.

Wichmann, F. A., Drewes, J., Rosas, P., & Gegenfurtner, K. R. (2010). Animal detection in natural scenes: Critical features revisited. Journal of Vision, 10 (4), 6–6.

Woolrich, M. W., Behrens, T. E., Beckmann, C. F., Jenkinson, M., & Smith, S. M. (2004). Multilevel linear modelling for fmri group analysis using bayesian inference. Neuroimage, 21 (4), 1732–1747.

Woolrich, M. W., Behrens, T. E., & Smith, S. M. (2004). Constrained linear basis sets for hrf modelling using variational bayes. NeuroImage, 21 (4), 1748–1761.

Woolrich, M. W., Ripley, B. D., Brady, M., & Smith, S. M. (2001). Temporal autocorrelation in univariate linear modeling of fmri data. Neuroimage, 14 (6), 1370–1386.

Yamashita, O., Sato, M.-a., Yoshioka, T., Tong, F., & Kamitani, Y. (2008). Sparse estimation automatically selects voxels relevant for the decoding of fmri activity patterns. NeuroImage, 42 (4), 1414–1429.

Ziv, Y., Burns, L. D., Cocker, E. D., Hamel, E. O., Ghosh, K. K., Kitch, L. J., . . . Schnitzer, M. J. (2013). Long-term dynamics of ca1 hippocampal place codes. Nature neuroscience, 16 (3), 264–266.

